# Dioecy in a wind-pollinated herb explained by disruptive selection on sex allocation

**DOI:** 10.1101/2025.01.07.631798

**Authors:** Kai-Hsiu Chen, John R. Pannell

**Author notes:** **Correspondance:** Kai-Hsiu Chen.

## Abstract

- The evolution of dioecy from hermaphroditism is widely thought to be a response to disruptive selection favouring males and females, driven by advantages of inbreeding avoidance, sexual specialization, or both. It has hitherto been difficult to uncouple the importance of these two hypotheses.
- We use phenotypes produced by experimental evolution to test the inbreeding avoidance hypothesis in populations from which sexual specialization can be effectively ruled out. We estimate the selfing rate and the shape of fitness gain curves under scenarios with and without inbreeding depression in experimental populations of wind-pollinated *Mercurialis annua* with high variation in sex allocation.
- We confirm a phenotypic trade-off between male and female allocation in *M. annua*. Individual selfing rates increased with pollen production. This dependence gave rise to strong disruptive selection on sex allocation due to its interaction with the mating system under the scenario of high inbreeding depression, especially for plants of medium and large sizes.
- Taken together, we demonstrate that inbreeding avoidance on its own can lead to disruptive selection on sex allocation, favouring the selection and maintenance of dioecy under wind pollination without associated benefits of sexual specialization.

## Introduction

Why hermaphrodites should ever evolve towards dioecy has long intrigued evolutionary biologists (Darwin, 1877; Charnov *et al*., 1976; Bawa, 1980; Thomson & Brunet, 1990; Renner & Ricklefs, 1995; Freeman *et al*., 1997; Ashman, 2006; Käfer *et al*., 2017). Two overarching explanations have been suggested. First, the ‘inbreeding avoidance’ hypothesis posits that unisexuality provides a fail-safe means of avoiding self-fertilization and the accompanying deleterious effects of inbreeding depression (Mather, 1940; Lewis, 1941; Charlesworth & Charlesworth, 1978). And second, the ‘sexual specialization’ hypothesis recognizes that the separation of sexes into different individuals allows males and females to express different trait values that optimize their respective fitness (Willson, 1979; Givnish, 1980), with the resulting sexual dimorphism resolving the sexual conflict and interference that may compromise fitness in simultaneous hermaphrodites (Abbott, 2011; Schärer *et al*., 2015). These two explanations for dioecy are not mutually exclusive: populations may initially evolve dioecy in response to selection for inbreeding avoidance but then subsequently evolve sexually dimorphic traits that confer on each of the two sexes benefits of specialization (Freeman *et al*., 1997; Charlesworth, 2018); or dioecy may evolve gradually from hermaphroditism via divergence in sex allocation and sexual specialization jointly (Lloyd, 1980a; Lesaffre *et al*., 2024b). However, it has not been possible to isolate the role of one or other of the two leading explanations for the evolution and maintenance of dioecy in any single species to date.

Progress in distinguishing between potential factors responsible for the evolution and maintenance of dioecy has been slowed by the difficulty to compare fitness among males, females, and hermaphrodites with a range of different sex allocation strategies in the same context. This difficulty arises because dioecious species typically comprise only males and females or, in hermaphroditic populations, the sex allocation of individuals is typically quite canalized. Moreover, while the males and/or females of some dioecious species show a degree of ‘leaky’ or inconstant sex expression (Delph & Wolf, 2005; Ehlers & Bataillon, 2007; Käfer *et al*., 2022), the sex allocation of these leaky phenotypes is typically very close to that of the corresponding ‘constant’ (or pure) males or females (Cossard & Pannell, 2019). We thus lack the sort of natural variation within populations that is needed for comparisons of the fitness of phenotypes across a wide range of alternative sex-allocation strategies. More formally, we lack estimates of the shape of ‘fitness gain curves’ on which much of the theory for the evolution of dioecy versus hermaphroditism is based (Charnov *et al*., 1976; Charlesworth & Charlesworth, 1981; Charnov, 1982; Charlesworth & Morgan, 1991; Campbell, 2000; Zhang, 2006; West, 2009; Fromhage & Kokko, 2010; Dorken & Van Drunen, 2018; Masaka & Takada, 2023; Lesaffre *et al*., 2024a).

Fitness gain curves relate the fitness of an individual in a population through its male and female components of reproductive success to its sex allocation, i.e., to the fraction of reproductive resources allocated to its male versus female functions (Charnov *et al*., 1976; Charnov, 1982; West, 2009). In the simplest terms, dioecy is predicted to be evolutionary stable when pure males and females have greater fitness than any intermediate (hermaphroditic) sex-allocation strategy, such that selection on sex allocation is disruptive and the curve relating the combined fitness through both male and female functions is u-shaped (Fig. **1**); such a scenario might be revealed, for example, by a positive second-order polynomial in a regression analysis (Lande & Arnold, 1983). In contrast, hermaphroditism should be stable to the invasion of sexual specialists (males or females, or individuals with highly male-biased or female-biased sex allocation) if selection is stabilizing and the fitness gain curve is n-shaped (potentially revealed by a negative second-order polynomial, for instance; Fig. **1**; Lande & Arnold, 1983). These ideas were applied to theory for the maintenance of dioecy versus hermaphroditism under the assumption of complete outcrossing of all phenotypes (e.g., hermaphrodites that are self-incompatible; e.g., Charnov et al. 1976), but also to the case where hermaphrodites are partially self-fertilizing, with the fitness of selfed progeny potentially diminished by inbreeding depression (e.g., Lloyd, 1975; Charlesworth & Charlesworth, 1978; Charlesworth, 1984 − the evolution of gynodioecy and androdioecy).

**Figure 1.**
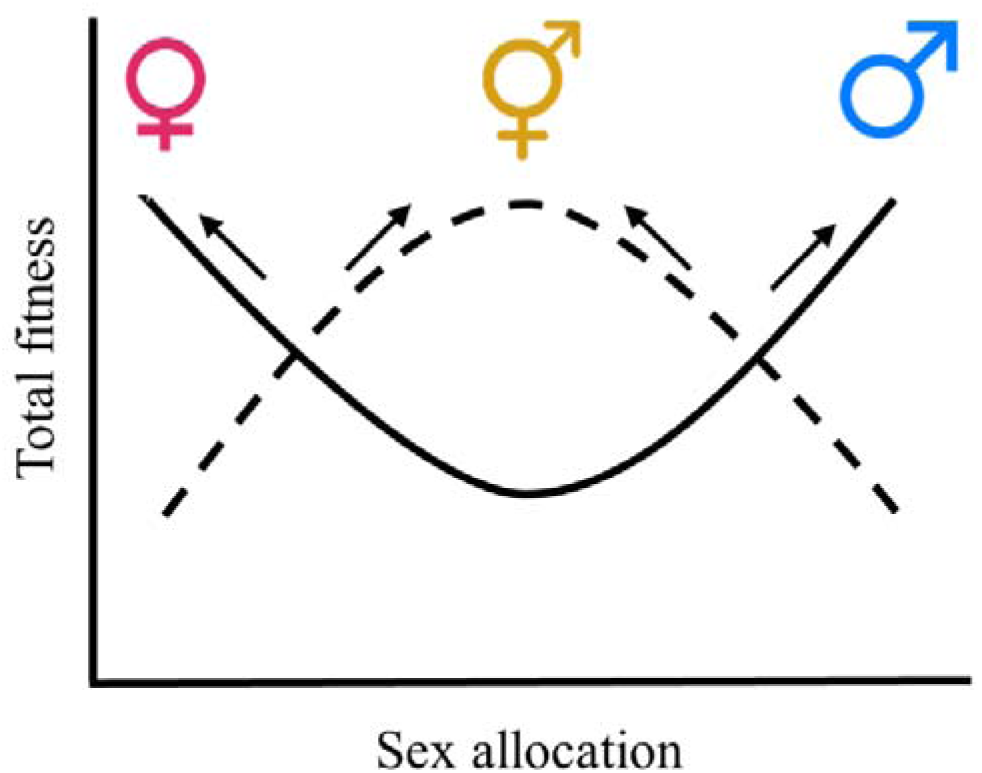
Conceptual diagram showing different selection schemes via total fitness on sex allocation (gender) within a population. Unisexual female and male individuals are favoured when the selection on sex allocation is disruptive (depicted by the solid line), rendering the population dioecious. In contrast, when the selection on sex allocation is stabilizing (depicted by the dashed line), hermaphroditism is favoured.

Models of the effect of selfing on the evolution of dioecy have most typically assumed a parametric fixed selfing rate independent of the distribution of sex allocation (Lloyd, 1975; Charlesworth & Charlesworth, 1978; Charlesworth, 1984), yet the selfing rate might often depend on floral strategies, including relative allocation to male and female functions (Denti & Schoen, 1988; Damgaard & Abbott, 1995; Chen & Pannell, 2024b). In particular, hermaphrodites allocating more to their male function might be expected to self-fertilize more of their seeds, either because pollinators of animal-pollinated species stay longer on plants with large floral displays (Karron *et al*., 2009; Christopher *et al*., 2021), or as a result of a simple ‘mass-action’ process (Gregorius *et al*., 1987; Holsinger, 1991), particularly in wind-pollinated species, whereby the selfing rate is determined by lottery competition.

The process of mass-action mating has important implications for the shapes of the male and female gain curves. In wind-pollinated species, allocating more resources to male function likely results in a relatively linear increase in male fitness through outcrossing, with increases in pollen production leading to corresponding increases in siring success (Charnov, 1982; Campbell, 2000; Aljiboury & Friedman, 2022). Yet we should also expect greater pollen production to increase the proportion of self-pollen in the pollen cloud around the plant’s stigmas, causing greater self-fertilization in self-compatible species (Gregorius *et al*., 1987; Denti & Schoen, 1988; Holsinger, 1991). While increased male allocation may therefore increase outcross siring success, it should also decrease female reproductive success in self-compatible species that suffer from strong inbreeding depression as a result of ovule or seed discounting (Lloyd, 1992; de Jong *et al*., 1999). Accordingly, when inbreeding depression is substantial, individuals allocating all their reproductive resources to either the male or female function should have higher fitness than those adopting a hermaphroditic strategy (Fig. **1**), leading to a u-shaped gain curve and the evolutionary stability of dioecy (de Jong *et al*., 1999; de Jong & Geritz, 2001).

The expectation of disruptive selection on sex allocation should also depend on the relative sizes of the plants concerned (de Jong *et al*., 1999; de Jong & Geritz, 2001). When selfing is increased by male allocation, disruptive selection should be strongest for the relatively larger individuals in a population, which produce absolutely more pollen. This is because larger individuals should have higher selfing rates for a given relative male allocation, with a steeper dependence of the selfing rate on sex allocation (Fig. **2**). In contrast, disruptive selection might be expected to be lowest for small individuals, which contribute less to the pollen cloud around their stigmas and thus self/fertilize less (Fig. **2**), allowing them to allocate to their male function with relative impunity. If so, small individuals might even experience stabilizing selection on their sex allocation in populations in which sex allocation in larger individuals otherwise favours the male and female extremes (de Jong & Geritz, 2001). In other words, we might expect disruptive selection in a population to dominate (because the large individuals contribute disproportionately to the next generation), even though small individuals may experience stabilizing selection. To our knowledge, this duality has not been modelled formally, but it would seem to be a likely scenario in wind-pollinated populations that conform to mass-action mating. While these ideas explicitly link sex-allocation theory with the inbreeding avoidance hypothesis for the evolution and maintenance of dioecy, they remain largely untested.

**Figure 2.**
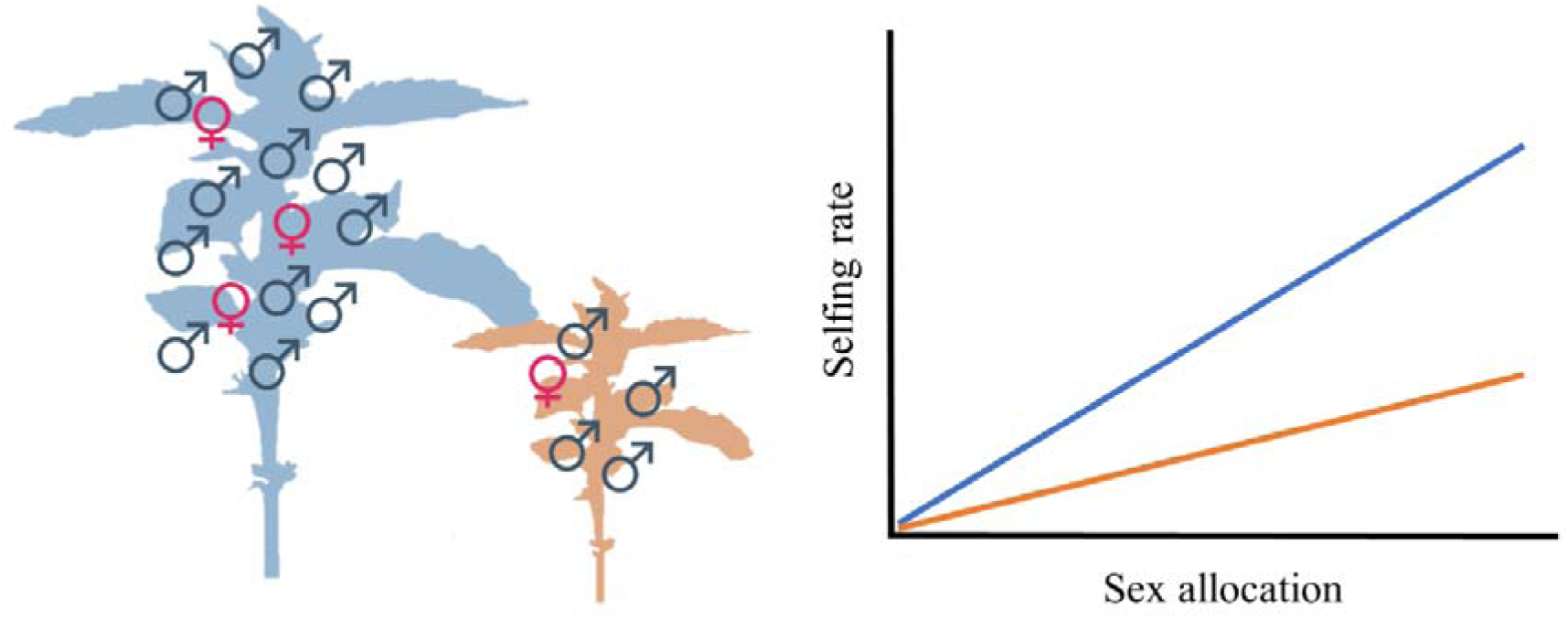
The expected effect of plant size on the selfing rates under the mass-action assumption (see main text for details). Given the same sex allocation (gender), a large plant (shown in blue) will produce absolutely more male flowers compared to a small plant (shown in orange). Thus, the expected selfing rate under the mass-action assumption, which is positively correlated to the male flower production, will be higher in the larger plant.

For empirical tests of theory for the stability of dioecy, we ultimately need to compare the fitness of individuals adopting a range of alternative sex-allocation strategies within a population, from fully male through hermaphroditism (ideally with a range of different sex allocations) to fully female (Fig. **1**). However, most plant species tend to be either dioecious or hermaphroditic, and species with intermediate sexual systems such as gynodioecy or androdioecy (where males or females, respectively, coexist with hermaphrodites) only allow comparisons between unisexuality and bisexuality for one of the sexual functions (Fritsch & Rieseberg, 1992; Weller & Sakai, 2005; Spigler & Ashman, 2012; Varga, 2021; Laugier *et al*., 2023). Although comparisons among related species with different sexual systems can provide valuable information (Steven & Waller, 2004; Sakai *et al*., 2006; Soza *et al*., 2012; Kwok & Dorken, 2022), they are often compromised by confounding trait variation and ecologies, and they do not replace comparisons among strategies within the same context. Comparisons among individuals expressing different sex-allocation strategies within the same population can be achieved by physical manipulation of the plants, e.g., through the removal of floral parts or flowers (Emms, 1993; Tomaszewski *et al*., 2018; Aljiboury & Friedman, 2022; Larue & Petit, 2024; Chen & Pannell, 2024a), but such manipulations are often difficult to achieve realistically and may not generate phenotypes that would occur in nature. The dearth of standing variation in sex allocation for testing patterns of selection within a population has therefore been a barrier to studies of the evolution and stability of separate sexes.

Here, we test the inbreeding avoidance hypothesis for the evolution of dioecy by studying the fitness of phenotypes of the self-compatible wind-pollinated dioecious annual plant *Mercurialis annua* across the full range of sex allocation, from male to female, and without confounding factors of sexual specialization (dimorphism). The phenotypes we used in our study were generated over the course of several generations of experimental evolution of females from initially dioecious populations from which males were removed; these females rapidly evolved substantial male allocation through the enhancement of a ‘leaky’ expression of male flowers (Cossard *et al*., 2021; Gerchen *et al*., 2024; Villamil *et al*., 2024). Critically, because males had been removed from these populations, they lacked the strong sexual dimorphism in vegetative traits that otherwise characterize natural populations of dioecious *M. annua* (see supplementary analyses in Table **S1**) and varied largely only in their sex allocation and, as our study demonstrates, in their selfing rates. Thus, although natural populations of dioecious *M. annua* are sexually dimorphic for several traits (Harris & Pannell, 2008; Tonnabel *et al*., 2019, 2022), which likely contributes to the evolutionary stability of dioecy, our model populations here allowed us to focus attention specifically on the inbreeding avoidance hypothesis.

We first confirmed the existence of a sex-allocation trade-off between male and female functions (Gerchen *et al*., 2024), the fundamental assumption of sex-allocation theory on which its predictions are based (Charnov, 1982; West, 2009). We then evaluated the merits of the inbreeding avoidance hypothesis by asking (1) how the selfing rate depends on sex allocation and individual plant size and (2) how total fitness depends on the selfing rate of individuals with different sizes and sex allocation strategies under contrasting scenarios of inbreeding depression. For each individual and its sex-allocation phenotype, we estimated male reproductive success on the basis of paternity assignment for a large sample of progeny. We estimated female reproductive success for all individuals in terms of a measure of seed production per individual for scenarios of contrasting inbreeding depression. With estimates of male, female, and total reproductive success, we finally inferred the shape of the respective fitness gain curves. Our study advances an understanding of the stability of dioecy to the invasion of alternative sex-allocation strategies on the basis of fitness comparisons over a range of physiologically realistic expressions of sex allocation that, to our knowledge, have hitherto not been realized.

## Materials and Methods

### Plant materials and experimental populations

*Mercurialis annua* (Euphorbiaceae) is a wind-pollinated annual herb widely distributed around the Mediterranean Basin and throughout central and western Europe (Tutin *et al*., 1976; Pannell *et al*., 2004). Natural diploid populations are dioecious, with individuals producing either female or male unisexual flowers, though leaky sex expression in both sexes (where individuals produce a small number of unisexual flowers of the opposite sex) is not uncommon (Cossard & Pannell, 2019, 2021; Villamil *et al*., 2022). To establish our experimental populations, we used seeds sampled from populations of females that had been evolving for eight generations in the absence of males (Cossard *et al*., 2021; Gerchen *et al*., 2024); while these females were phenotypically similar to those found in natural populations of diploid *M. annua* (see Table **S1** for descriptions of the ancillary traits), they had evolved substantially enhanced ‘leaky’ production of male flowers that resemble in important respects monoecious individuals from polyploid individuals (Pannell *et al*., 2008), but with substantially greater variation in male-flower production.

Seeds for our study were sown in seedling trays in July 2022 and grown for five weeks in the greenhouse of the University of Lausanne, Switzerland (Cossard *et al*., 2021; Gerchen *et al*., 2024; Method **S1**). In mid-August, we set up three common-garden populations as independent replicates in hexagon-shaped plots, each comprising 61 individuals (Method **S1**). The populations were set up at least 150 m apart from each other on the University campus, minimizing gene flow among them (the average outcrossing mating distance within the populations was 28.5 cm; Table **S2**). The seedlings were potted individually in 16 cm pots with soil (Ricoter substrate 140) and slow-release fertilizer (Hauert Tardit 6M pellets; 5 g fertilizer L–1 of soil) and were automatically watered throughout the period of the experiment of around 14 weeks. We harvested the above-ground parts of the plants and measured their sex allocation and other traits in mid-October when the night temperature dropped below 5 °C and the plants ceased growing.

### Phenotyping of sex allocation and biomass

To quantify the sex allocation of each individual, we counted the number of female (N_F_) andmale (N_M_) flowers produced on the whole plant by detailed inspection throughout all the branches at the time of harvest. For each individual *i*, we further calculated its functional gender (*G*, hereafter gender) (*sensu* Lloyd, 1980) in terms of maleness as G_i_ = N_M,i_/(Ex N_F,i_ + N_M,i_), where the equivalence factor, *E*, is the ratio of the number of total male flowers to total female flowers in the respective population. The inclusion of *E* in the formula not only guarantees that the mean gender of the population is 0.5, reflecting the fact that exactly half of all genes passed to progeny are via male and the other half via female function, but it also allows measuring male and female allocation in any units (see Harris & Pannell (2008) for an example of estimates of allocation in dioecious *M. annua* in terms of biomass and nitrogen content). After the phenotyping, we kept all the harvested parts in bread bags to dry at room temperature for three weeks. We then measured their biomass and gathered and counted mature seeds that had been dispersed from capsules into the bags.

### Paternity analysis and the estimate of the selfing rate

To estimate the selfing rate of each individual, we used variation at nine microsatellite markers to assign paternity to mature seeds (modified from Machado et al., 2017). Leaf samples of the parental individuals were collected upon harvest and dried in silica gel prior toDNA extraction. We also extracted DNA from a sample of progeny produced by each of them. Specifically, we randomly sampled five to ten mature seeds from the total seed family of each female parent, with five, six, seven, eight, nine, and ten seeds sampled for individuals with five to 100, 101 to 200, 201 to 300, 300 to 400, 400 to 500, and > 500 seeds, respectively; in the few cases in which individual produced fewer than five seeds, we sampled all of them. In total, we extracted DNA from 181 parents and 958 seed progeny. We could assign paternity to 915 of the ∼13,800 seed progeny in the population, which amounted to a sampling effort of 6.6%. Total DNA was extracted from the leaves and seed samples using the BioSprint 96 DNA Plant Kit (Qiagen, Germany) according to the manufacturer’s instructions and eluted in 100 µl of distilled water.

PCR amplification was carried out in a final volume of 10 µl, including 5 µl of 2× Multiplex PCR Master Mix (Qiagen, Germany), 0.2 µl of diluted DNA, 2.8 µl of distilled water, and 2.0 µl of multiplex containing variable primer concentrations. The nine microsatellite markers were grouped into two multiplexes modified from Machado et al. 2017. Thermal cycling was performed in a TProfessional Standard Thermocycler (Biometra GmbH, Göttingen, Germany) as follows: 95 °C for 15 min; 33 cycles at a temperature of 94 °C for 30 s, 90 °C for 90 s, and 72 °C for 90 s; and a final step at 72 °C for 10 min before cooling down to 4°C. PCR products were analyzed by capillary electrophoresis on an ABI3100 Genetic Analyzer (Applied Biosystems), with an internal size standard GeneScan-500 LIZ. Fragment length analyses and scoring were performed with GeneMapper v 6.0 (Applied Biosystems).

Paternity analyses were conducted for the three populations separately to assign the most likely father to each seed for which more than five loci were genotyped. Here, we used the software Cervus v 3.0.7, assuming a relaxed confidence level of 80% and an error rate of 0.01 (Kalinowski *et al*., 2007). *M. annua* is self-compatible, and the female selfing rate (hereafter selfing rate) of each individual was estimated by the proportion of selfed seeds to the number of total seeds whose paternity was successfully assigned.

### Calculation of female, male, and total fitness

The annual life cycle of *M. annua* allows us explicitly to estimate lifetime fitness in terms of the number of seeds produced and sired by each individual. In this study, following the practice of most empirical studies (e.g., Karron & Mitchell, 2012; Briscoe Runquist *et al*., 2017; Chen & Pannell, 2023; Hou *et al*., 2024), we attributed the fitness gained via selfed progenies equally to female and male functions rather than attributing twice the fitness of selfed progeny to only the female function, as is typical in theoretical studies (e.g., Charlesworth & Charlesworth, 1987b; Lesaffre *et al*., 2024a). However, note that inferences for total fitness based on the two contrasting approaches are entirely equivalent. Fitness components were estimated under two scenarios using selfing rates (*S*) estimated from the paternity analysis: one in which inbreeding depression (δ) was assumed to be zero; and one in which δ was assumed to equal one. The actual level of inbreeding depression in wild populations of diploid *M. annua* remains unknown, with a previous study showing a negligible level of inbreeding depression estimated in hexaploid dioecious populations (Eppley & Pannell, 2009; Pujol *et al*., 2009), and another unpublished study showing considerable inbreeding depression in diploid populations under experimental evolution (E. Le Faou and J.R. Pannell, unpublished data). The scenarios assumed in our analysis reflect the two extreme levels of inbreeding depression, allowing us to explicitly evaluate the effects of inbreeding depression on the patterns of selection on sex allocation. (See Fig. **S2** for supplementary analyses assessing the threshold values of inbreeding depression relevant to selection on sex allocation in plants of different sizes.)

We calculated the female fitness of each individual *i* as w_F,i_ = N_T,i_ x [S_i_(1-o) + (1-S_i_)], where S_i_ is the estimated selfing rate of individual *i*. Under the scenario of δ = 0, where selfing causes no inbreeding depression, female fitness is just the number of mature seeds of the individual (N_T,i_). When δ = 1, selfed progenies do not contribute to the next generation at all.

We calculated male fitness of each individual *i* using the paternity share of each dame derived from the paternity analysis as 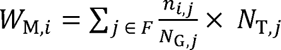, where n is the number of seeds sired by individual *i* on individual *j*, N_G,j_ is the number of genotyped seeds of dame *j*, and N_T,j_ is the total number of mature seeds of dame *j*. For selfed progeny (for which *i* = *j*), fitness accrued was discounted by a factor δ. We calculated total fitness of individual *i* as w_T,i_ = W_F,i_+ w_M,i_ for each of the two scenarios of inbreeding depression.

### Statistical analysis

We conducted all the analyses within the *R* statistical framework v 4.0.3 (R Core Team, 2021). We checked the fit of the models with the package *DHARMa* (Hartig, 2019) and QQ plots. The detailed structures of each regression model can be found in the Supporting Information (Method **S2**). The general effects of the explanatory variables in each model were extracted using likelihood ratio tests using the *drop1* function.

To evaluate the trade-off between investment in female versus male flower numbers (revealed by a negative coefficient), we used a zero-inflated generalized linear mixed model (*glmmTMB* function in package g*lmmTMB*; Brooks et al., 2017), setting the number of female flowers (non-negative integer numbers) as the response variable with a negative binomial distribution. Male flower number, plant size, and population were set as the explanatory variables with two-way and three-way interaction terms. For model convergence, the male flower number was standardized to a mean of zero and a standard deviation of one for each population.

Above-ground biomass was log-transformed (hereafter referred to as ‘plant size’). Note that plant size was analysed as a continuous variable in all analyses. To examine the interactive effect with plant size, we extracted the predicted coefficients at three levels of plant size (small, medium, and large), with size thresholds defined as plants of mean size minus SD (at the 16% quantile), mean size (at the 50% quantile), and mean size plus SD (at the 84% quantile), respectively (Fig. **S3**).

To investigate how the selfing rate depends on sex allocation (i.e., gender) and the size of focal plants in the three populations, we constructed a generalized linear mixed model (*glmer* function in package *lme4*, Bates et al. 2015) using the selfing rate of each mother individual as a binomial response variable. Gender, plant size, and population were set as the explanatory variables with two-way and three-way interaction terms. We set individual identity as a random variable to account for non-independence of seeds of the same individual. To assess how inbreeding depression affects the dependency of different components of fitness on gender and plant size under two inbreeding depression scenarios, we fitted relative female, male, and total fitness as the response variable in three separate linear mixed models (*lmer* function in package *lme4*, Bates et al. 2015).

The relative fitness of each individual was calculated by dividing the fitness by the mean of the focal population (Lande & Arnold, 1983). To detect the non-linear dependence of fitness on gender in each model, we set both linear and quadratic terms of gender as the explanatory variables accompanied by interactions with plant size, scenarios of inbreeding depression, and population. We weighted the variances by plant size to indicate that plants of different sizes have different variances (larger plants have larger variance in fitness; see Supporting Information Methods **S2**). We set the identity of parental individuals as a random variable to account for the fact that the fitness estimates under the two inbreeding depression scenarios were not independent (i.e., the same individual). We first extracted *P* values of the general effects of the explanatory factors via likelihood tests using the *drop1* function. We further used the *emtrends* function in the *emmeans* package to extract from the fitted models and conducted post-hoc comparisons of the linear and quadratic coefficients and their standard errors (Lenth, 2020), again for both scenarios of contrasting inbreeding depression and for different plant sizes.

## Results

### Variation in sex allocation

Plants in the experimental population varied greatly in their size and their male and female reproductive allocations, ranging along a continuum from pure females to pure males (Fig. **3a**; Fig. **S1**). Biomass and sex allocation were independent (*r* = 0.04, *P* > 0.05) and showed no difference among populations (Table **S2**). Individuals produced an average of 238 ± 282 and 316 ± 449 female and male flowers, respectively (mean ± SD; *N* = 180; see Table **S2** for each population; Fig. **3a**). Larger plants produced more female and male flowers (*P* < 0.001). Our data also revealed a clear sex-allocation trade-off, with a negative dependence of female flower number on male flower number (*P* < 0.001) and the strength of the trade-off being greater for larger plants (Fig. **3b** and Table **S3**). See Fig. **S4** for an evaluation of the non-linear sex-allocation trade-off.

**Figure 3.**
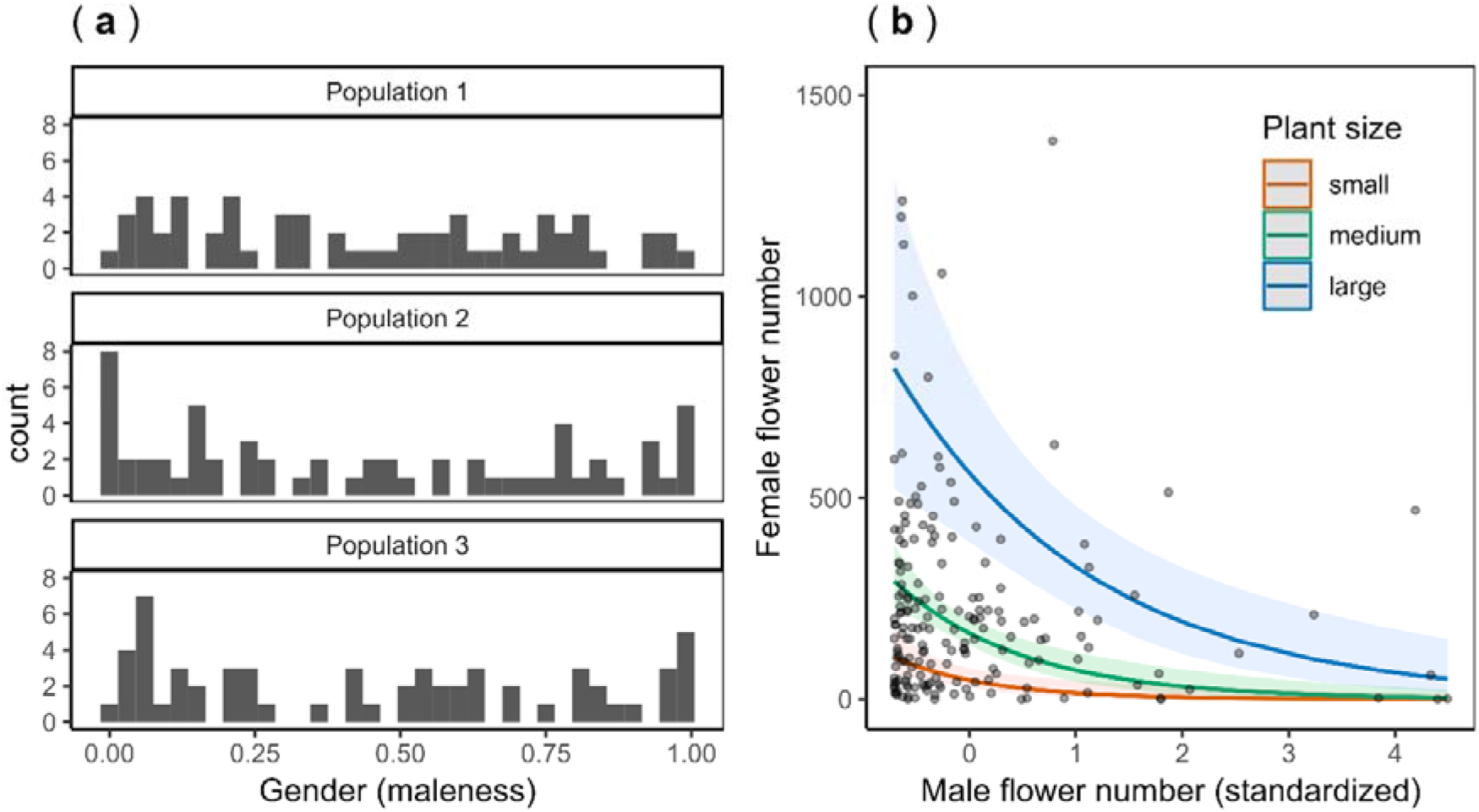
Plots showing the distribution of gender phenotypes in the studied populations (**a**) and the trade-off between female and male flower numbers of the individuals (**b**) of *Mercurialis annua*. In (**a**), a value of zero in gender indicates an individual with only female flowers, whereas a value of one indicates an individual with only male flowers. In (**b**), the trade-off lines for plants of different sizes were estimated by the generalized linear mixed model with the shaded ribbons indicating the 95% confidence interval of the corresponding regression lines. Hereafter to present the interactive effect with plant size, the regression lines for three levels of size (small, medium, and large) were shown, reflecting plants of mean size minus SD, mean size, and mean size plus SD, respectively. Note that one raw data point with an extreme number of female flowers (2108 female flowers) was not shown to avoid compression of the y-axis (see also **Fig.** S1).

### Dependence of self-fertilization on gender

In total, 948 seeds were successfully genotyped for at least five loci, for which paternity was assigned to a single father for 915 seeds (see also Table **S2** for details of each population).

Overall, the average selfing rate was 29.6% (mean over *N* = 172 individuals; Table **S2**). The selfing rate was higher for individuals with greater male allocation (and greater male gender) (*P* < 0.001; Fig. **4**), with the dependence tending to be steeper for larger plants (Fig. **4**), although there was no significant interaction among gender, size, and population (Table **S4**). See also Fig. **S5** for the positive dependence of the selfing rate on absolute male allocation, i.e., male flower number.

**Figure 4.**
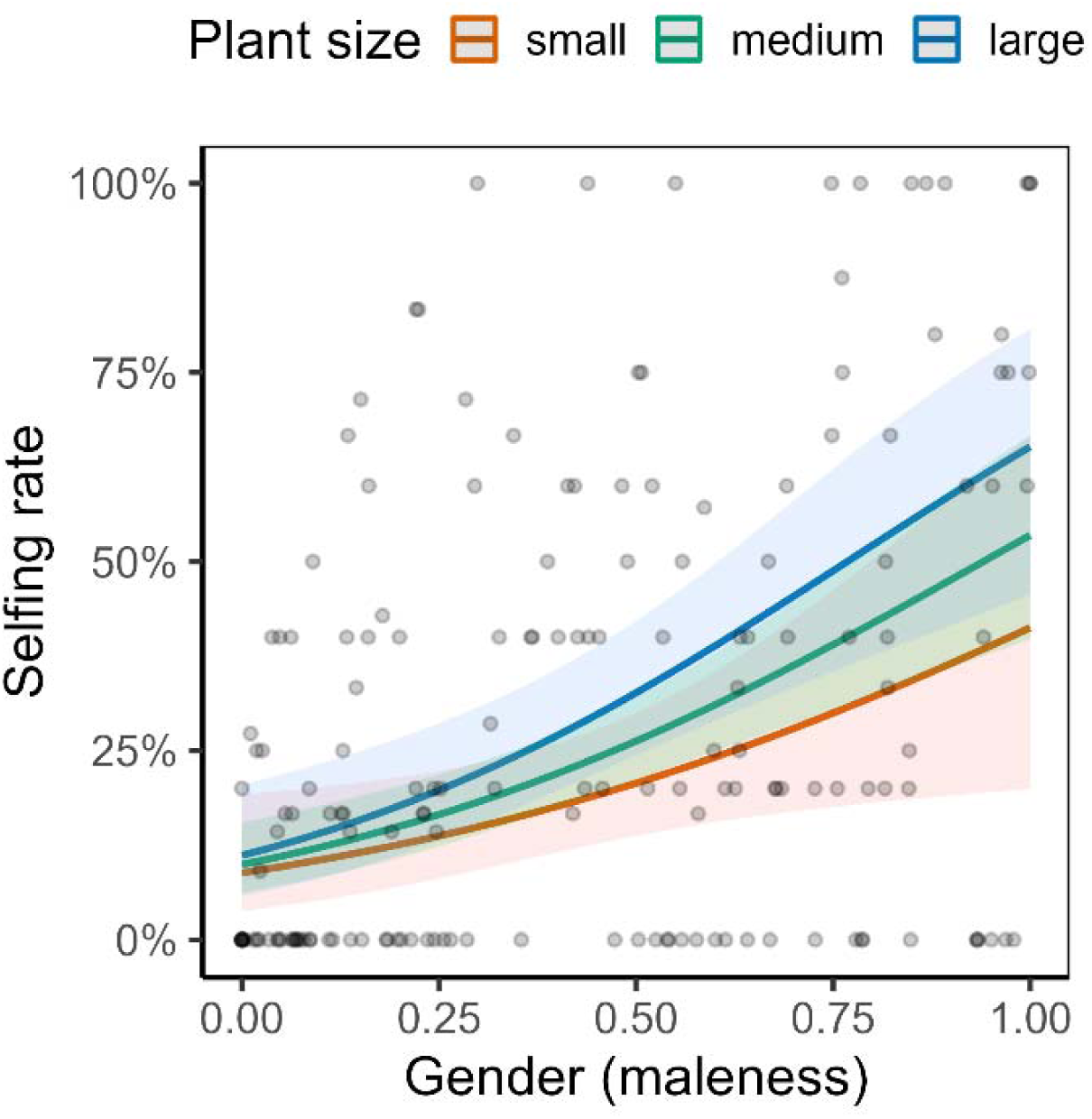
Plot showing the effect of gender (maleness) on the selfing rate of studied individuals of *Mercurialis annua* (*N* = 172, individuals producing no mature seed were not included). Note that although the interaction with plant size was not significant, the slope trended to be steeper in larger plants. The shaded ribbon indicates the 95% confidence interval of the regression lines. Note that there was one individual with a gender of zero that had a selfing rate > 0. This inference is likely the result of sampling error during the phenotyping, where we might have overlooked male flowers on that individual.

### Dependence of fitness on gender, plant size, and inbreeding depression

Patterns of mating and fitness tended to be similar across populations: because all higher-order interaction terms involving population were non-significant (*P* > 0.05; Table **S5**), we dropped interactions with population in presenting the results (Table **1**; see also Fig. **S6** and **S7** for a comparison of selection and effect sizes among populations). Table **1** presents the linear and quadratic coefficients of different components of fitness with gender under the two inbreeding depression scenarios for plants of different sizes.

**Table 1.** Comparisons of linear and quadratic coefficients of relative female (**a**), male (**b**), and total (**c**) fitness on gender between the two scenarios of inbreeding depression (δ) in plants of different sizes. Standard errors of the estimates of coefficients are indicated in the parenthesis. A significant comparison indicates that the coefficients under the two scenarios are statistically different, based on post-hoc pairwise comparisons (see details in the main text). Significant coefficients and comparisons are noted by asteroids. Notes: ∗ P < 0.05, ∗∗ P < 0.01, ∗∗∗ P < 0.001.

| | $\delta = 0$ | $\delta = 1$ | Comparison |
| --- | --- | --- | --- |
| (a) Relative female fitness |  |  |  |
| Small plants | Linear: 0.87 (1.86) | Linear: 0.33 (1.86) |  |
|  | Quadratic: -1.03 (1.91) | Quadratic: -0.54 (1.91) |  |
| Medium plants | Linear: -2.01 (1.09) | Linear: -3.92 (1.09) *** | *** |
|  | Quadratic: 0.35 (1.1) | Quadratic: 1.97 (1.1) | *** |
| Large plants | Linear: -4.92 (1.75) ** | Linear: -8.21 (1.75) *** | *** |
|  | Quadratic: 1.74 (1.77) | Quadratic: 4.51 (1.77) * | *** |
| (b) Relative male fitness |  |  |  |
| Small plants | Linear: 1.58 (1.78) | Linear: 1.28 (1.78) |  |
|  | Quadratic: -1.54 (1.82) | Quadratic: -1.07 (1.82) |  |
| Medium plants | Linear: 0.92 (1.05) | Linear: 0.08 (1.05) | * |
|  | Quadratic: 0.43 (1.06) | Quadratic: 2.06 (1.06) | *** |
| Large plants | Linear: 0.27 (1.68) | Linear: -1.11 (1.68) |  |
|  | Quadratic: 2.4 (1.71) | Quadratic: 5.19 (1.71) ** | *** |
| (c) Relative total fitness |  |  |  |
| Small plants | Linear: 1.23 (1.32) | Linear: 0.81 (1.32) |  |
|  | Quadratic: -1.30 (1.35) | Quadratic: -0.83 (1.35) |  |
| Medium plants | Linear: -0.55 (0.77) | Linear: -1.92 (0.77) * | *** |
|  | Quadratic: 0.38 (0.78) | Quadratic: 2 (0.78) * | *** |
| Large plants | Linear: -2.35 (1.24) | Linear: -4.68 (1.24) *** | *** |
|  | Quadratic: 2.07 (1.26) | Quadratic: 4.86 (1.26) *** | *** |

The relationship between relative fitness and gender (maleness) depended strongly on plant size and scenarios of inbreeding depression, for both female function (*P* < 0.001 and < 0.01 for three-way interaction terms involving linear and quadratic terms of gender, respectively; Table **S5**; Fig. **5a**) and male function (*P* > 0.05 and < 0.01 for three-way interaction terms involving linear and quadratic terms of gender, respectively; Table **S5**; Fig. **5b**). For example, in small plants, relative male and female fitness components did not depend on gender for either scenarios of inbreeding depression (*P* > 0.05 for all coefficients; Table **1**; Fig. **5a** and **b**), whereas in large plants, relative female fitness negatively and convexly depended on gender for a scenario assuming δ = 1 (for both sexual functions, *P* < 0.001 and < 0.05 for linear and quadratic terms respectively; Table **1**; Fig. **5b**), but it depended only linearly on gender for a scenario assuming δ = 0 (for both sexual functions, *P* < 0.01 and > 0.05 for linear and quadratic terms respectively; Table **1**; Fig. **5b**).

**Figure 5.**
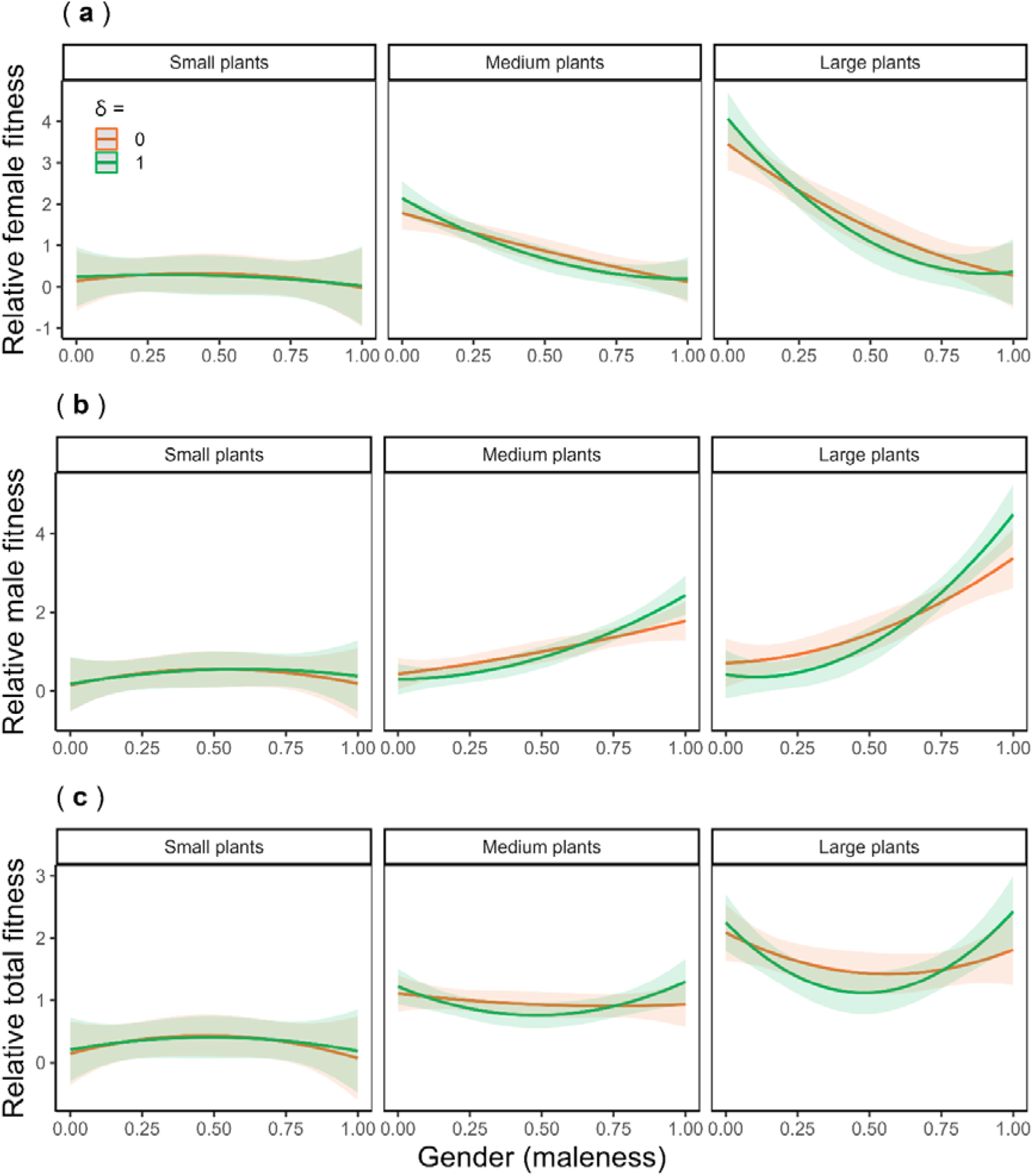
Plots showing the interactive effects of plant size, degree of inbreeding depression, and gender on relative female (**a**), male (**b**), and total (**c**) fitness of *Mercurialis annua*. Fitness was estimated under two scenarios of inbreeding depression (δ) of zero and one, depicted by orange and green lines, respectively. The shaded ribbons indicate the 95% confidence interval of the corresponding regression lines. The interactions with population were not significant thus the interactive effects with population were not shown (see Table **1** and Table **S4** for the *P* values).

When considering total fitness under different inbreeding depression scenarios, patterns of selection on gender were different for plants of different sizes (Table **S5**). Relative total fitness depended disruptively on gender in large plants under a scenario of δ = 1 (*P* < 0.001 and < 0.001 for linear and quadratic terms respectively; Table **1**; Fig. **5c**), whereas the disruptive dependence was non-significant when assuming δ = 0 (*P* > 0.05 for both terms; Table **1**; Fig. **5c**). In contrast, there was no detectable selection on gender in small plants regardless of the scenarios of inbreeding depression (Table **1**; Fig. **5c**)

## Discussion

The wide phenotypic variation in sex allocation displayed in our experimental populations of wind-pollinated *M. annua* allowed us to evaluate the dependence of the selfing rate and total plant fitness on allocation to male and female functions. Three results stand out. First, we confirmed the existence of a clear trade-off between male and female allocation (Gerchen *et al*., 2024), as assumed in theories of sex-allocation and life-history evolution (Charnov, 1982; West, 2009). Second, the selfing rate of individuals depended positively on male allocation and greatly altered the shape of female and male fitness gain curves under scenarios of strong inbreeding depression, especially for plants of medium and large sizes. And third, sexual interference caused by the interaction between sex allocation and the mating system led to consistent patterns of strong disruptive selection on sex allocation, as required for the evolution and maintenance of dioecy (de Jong *et al*., 1999; de Jong & Geritz, 2001).

### Non-linear trade-off in allocation between female and male function

We found a constant negative association between the number of male and the number of female flowers for each size class in the population. This result provides evidence for a sex-allocation trade-off for plants that likely had a similar resource status, as assumed by sex-allocation theory (Charnov, 1982; West, 2009). In dioecious species, the production of sons versus daughters is usually a ‘zero-sum game’, such that sex-allocation trade-offs are almost axiomatic. However, it has been difficult to demonstrate such trade-offs in hermaphroditic species, likely because critical covariates of sex allocation such as resource status have been overlooked, or because male and female functions draw on different resources (de Jong, 1993; Campbell, 2000; Johnson & Nassrullah, 2024; summarized in Ashman, 2003; Mazer et al., 2007). A negative correlation between male and female functions has been found in some monoecious species that produce a mixture of unisexual flowers, e.g., *Astilbe biternata* (Olson & Antonovics, 2000), *Pinus silvestris* (Savolainen *et al*., 1993), and *Zea mays* (Garnier *et al*., 1993), but see *Begonia semiovata* (Agren & Schemske, 1995). Our confirmation of a trade-off between the production of male and female flowers in *M. annua* under experimental evolution (Gerchen *et al*., 2024) adds to this evidence.

Importantly, the sex-allocation trade-off revealed for *M. annua* deviates strongly from the linear relation assumed in sex-allocation theory (e.g., Charlesworth and Charlesworth, 1981; Charnov, 1982), with its shape being significantly concave (see also Fig. **S1** for a supplementary analysis). This concavity would appear to reflect advantages of the ‘economics of scale’ for each sexual function and thus an advantage of specialization (Reekie & Avila-Sakar, 2005; Saeki *et al*., 2014), possibly linked to negative physiological interference between the sexual functions of individuals producing both male and female flowers, e.g., brought about by hormone regulation, gene expression, and nutrient acquisition (Diggle *et al*., 2011; Golenberg & West, 2013; Sobral *et al*., 2016; Jabbour *et al*., 2022). Sex expression in *M. annua* is regulated by phytohormones at the early stage of floral development, with auxin and cytokinin inducing male and female floral buds, respectively (Louis *et al*., 1990). It is thus plausible that the production of a mixture of male and female flowers by monoecious individuals may partially disrupt a finely tuned regulatory network required for the production of either male or female flowers (Durand & Durand, 1991; Golenberg & West, 2013). If so, such physiological interference would reduce the efficacy in flower production for individuals with intermediate allocation, at least in our study populations that have evolved under experimental conditions, and it may generally contribute to ecological advantages of sexual specialization in allocation in wild plant populations. The specialization in sex allocation, as implied by the nonlinear resource trade-off, likely favours dioecy in *M. annua* irrespective of the effects of inbreeding, because selection on sex allocation is disruptive for large plants even in the absence of inbreeding depression.

### Effect of selfing and inbreeding depression on female fitness

The selfing rate of individuals in our experimental populations increased with their relative and absolute allocation to male function. A positive dependence of selfing on male allocation has been demonstrated within a population for several insect-pollinated hermaphroditic species, both at the individual and flower levels (Damgaard & Abbott, 1995; Harder & Barrett, 1995; Karron *et al*., 2004; Williams, 2007; Chen & Pannell, 2024b), but it has hitherto not been reported for any wind-pollinated angiosperm (though such a pattern has been reported for a gymnosperm; Denti & Schoen, 1988). Although many wind-pollinated angiosperms are self-incompatible or dioecious (Renner & Ricklefs, 1995; Vogler & Kalisz, 2001; Friedman & Barrett, 2008), the selfing rate should be sensitive to male allocation in self-compatible wind-pollinated plants. Our study suggests that the potentially monoecious or hermaphroditic precursors of currently dioecious wind-pollinated species, which will often have been self-compatible (Charlesworth, 1985), may well have experienced the negative effects of self-fertilization, and that these might have contributed to selection for unisexuality.

The effect of sex allocation on the selfing rate also depended on the size of the plants considered. This finding corresponds to the predictions of the ‘mass-action’ model (Gregorius *et al*., 1987; Holsinger, 1991), in which absolute allocation to male function increases the proportion of self-pollen in the local pollen cloud and thus increases the selfing rate of the seeds produced (see also Fig. **S5**). The positive dependence of selfing on sex allocation was steeper in larger plants, because larger plants produced more male flowers in absolute terms than smaller plants. In contrast, male flowers produced by small plants had a much milder effect on the selfing rate because of their smaller resource budget. As a result, most of the ovules produced by the small plants, regardless of their male allocation, were outcrossed by pollen produced by the larger neighbouring individuals.

We found that the effect of male allocation on the rate of self-fertilization in *M. annua* causes a dependence of female fitness on sex allocation under a scenario of high inbreeding depression, especially in medium- and large-sized plants. When inbreeding depression was assumed to be zero in our analyses, female fitness depended linearly and positively on the allocation to female function, implying a mostly linear female gain curve. Under high inbreeding depression, by contrast, an elevated level of ovule discounting in individuals with increased male allocation should impose substantial fitness costs on female function. As a result, females that avoid allocating to their male function should enjoy higher relative female fitness, leading to an accelerating female gain curve (de Jong *et al*., 1999). Our results thus provide empirical evidence for the joint effect of the mating system and inbreeding depression on the female fitness gain curves due to sexual interference, as modelled by Charlesworth & Charlesworth (1981), de Jong *et al*. (1999), and Lesaffre *et al*. (2024a); see Chen & Pannell (2024b) for another recent example in an insect-pollinated species.

### Implications for the evolution of sexual systems in wind-pollinated plants

We found that male fitness in *M. annua* did not level off with an increased allocation to male function, indicating a linear (non-saturating) male fitness gain curve. Our estimates of male fitness when inbreeding depression was assumed to be zero support the general view that wind-pollinated species should have a linear male gain curve, because wind is not easily saturated with pollen in the same way that an insect’s body may be (Charnov, 1982; Lloyd & Bawa, 1984). The inferred linear male fitness gain curve for *M. annua* is thus similar to those for the wind-pollinated self-incompatible *Ambrosia artemisiifolia* (Nakahara *et al*., 2018; Aljiboury & Friedman, 2022) and self-compatible *Picea glauca* (Schoen & Stewart, 1986).

Our joint estimates of female and male fitness clearly indicate that dioecious sexual systems should be favoured as an evolutionary response to promote male fitness and avoid self-fertilization in self-compatible wind-pollinated species (de Jong *et al*., 1999). Considering total fitness gained via the two sex functions under strong inbreeding depression, our study points to the action of strong disruptive selection on sex allocation in medium- and large-sized plants of *M. annua*, favouring greater separation of the sexes in all three replicate populations (Fig. **S6**). Although an accelerating female gain curve due to ovule discounting may be common in both insect- and wind-pollinated species (Karron *et al*., 2004; Williams, 2007; Chen & Pannell, 2024b), the male gain curve is likely to be saturating in insect-pollinated species (de Jong & Klinkhamer, 1994), potentially precluding disruptive selection on sex allocation and rendering the evolutionary stable sexual system hermaphroditic or gynodioecious (de Jong *et al*., 1999; de Jong & Geritz, 2001). Our finding of disruptive selection on sex allocation in *M. annua* may thus help to explain the common association between dioecy and wind pollination in flowering plants (Renner & Ricklefs, 1995; Vamosi *et al*., 2003; Friedman & Barrett, 2009) and the frequent evolution of dioecy from hermaphroditism following shifts to wind-pollination, e.g., in *Fraxinus* (Wallander, 2008), *Schiedea* (Weller *et al*., 1995), and *Thalictrum* (Soza *et al*., 2012).

We also found that disruptive selection on sex allocation was particularly strong for large plants and almost absent for small plants (see also Figure **S2**). This result would thus seem to imply that size-dependent sex allocation ought to be selected in *M. annua*, with the large individuals being strictly unisexual and small individuals expressing a wider range of sex allocation (de Jong *et al*., 1999; de Jong & Geritz, 2001). The fact that wild individuals of *M. annua* are in fact predominantly unisexual, irrespective of their size, may indicate that dioecy has been stabilized largely by the selection on mid- and large-sized individuals, which produce almost all their parents’ descendants: the smallest 16% of plants (i.e., plants with a size smaller than one standard deviation from the mean size) produced on average only 18.7 ±18.3 seeds, compared with the average seed production of 106 ± 159.8 in the populations as a whole), and it is thus plausible that they adopt a unisexual strategy because selection on their sex allocation is correspondingly weak.

### Concluding remarks

Selection for inbreeding avoidance or selection for traits that promote advantages of sexual specialization have long been identified as the two most likely mechanisms favouring the evolution of dioecy in plants (Bawa, 1980; Thomson & Brunet, 1990; Freeman *et al*., 1997; Pannell & Jordan, 2022). Although separate sexes in wild populations of dioecious *M. annua* are likely favoured by advantages of sexual specialization not linked to inbreeding depression (Eppley & Pannell, 2007; Tonnabel *et al*., 2019), our results now indicate that disruptive selection on sex allocation, and thus the potential evolution of separate sexes, can arise in populations through the avoidance of the negative effects of selfing when the male fitness gain curve is not too saturating, even in the absence of advantages of sexual dimorphism.

Thus, while sexual specialization on ancillary traits may reinforce selection for separate sexes and the evolution of dioecy with sexual dimorphism (Lesaffre *et al*., 2024b), our study demonstrates that inbreeding avoidance on its own can drive the evolution of dioecy (see Fig. **S8** for the negligible effects of ancillary traits on fitness from supplementary analyses). Our study thus provides support for fundamental elements of theory of sex allocation that have been difficult to test or validate for lack of suitable phenotypic variation.

## Acknowledgements

We thank D. Savova Bianchi, A. Hoang, E. Bochaton, L. Wagneur, and L. Cordella for data collection and G. Cossard, X. Li, J. Gerchen, N. Villamil-Buenrostro, E. Le Faou, and their teams of helpers for maintaining the selection experiment over the years. JRP acknowledges the Swiss National Science Foundation for ongoing funding of his research on the evolution of plant sexual systems (SNSF grant 310030_185196).

## Author contributions

KHC and JRP designed the project; KHC collected the data; KHC analysed the data with input from JRP; both authors wrote and edited the manuscript.

## Data availability

The data that support the findings of this study is openly available in Zenodo (10.5281/zenodo.14542261).

## Supporting Information

**Method S1.**
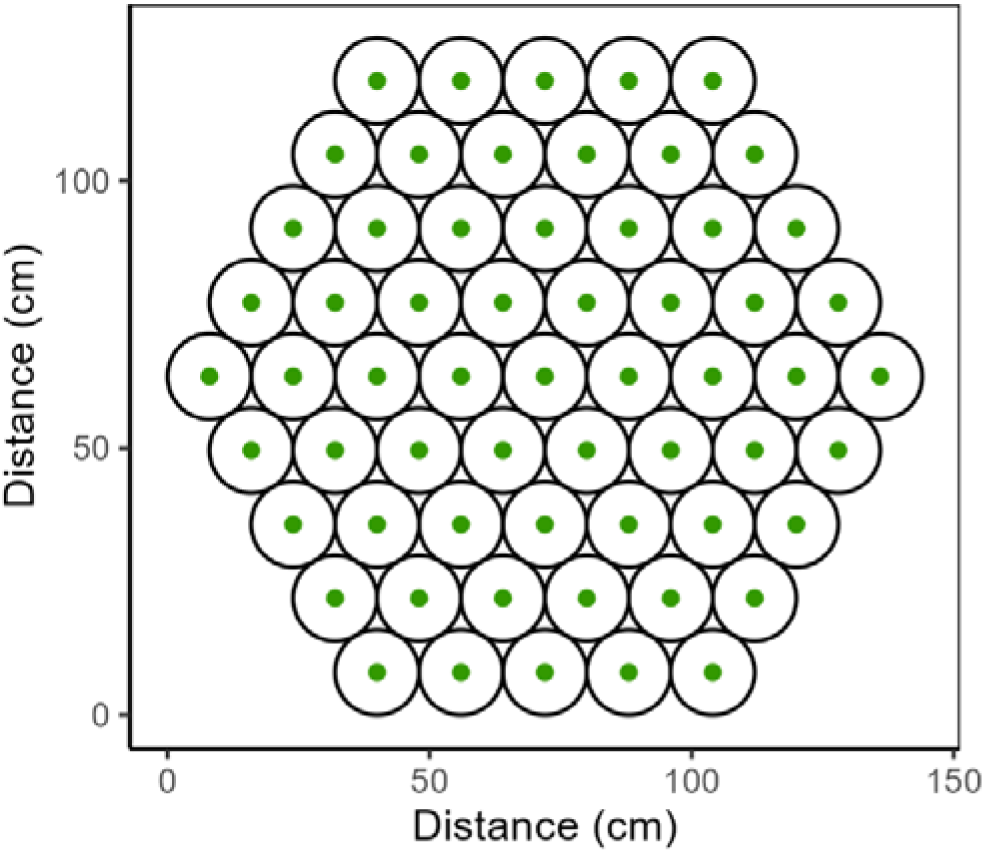
Around 225 seeds of *Mercurialis annua* were randomly sampled from the bulk-harvested seed pools of each of the three replicate populations of an ongoing experiment in which females had evolved substantial male-flower production after the experimental removal of males (Cossard et al., 2021; Gerchen et al., 2024). The seeds had been stored since their harvest in 2020 at 4 °C. The sampled seeds were sown individually in 9 x 5 well trays in a greenhouse at the University of Lausanne in July 2022. After five weeks, 61 seedlings (green dots in the figure below) were randomly selected and replotted into pots with a diameter of 16 cm (empty circles in the figure below) and were arranged in a five-layered hexagon for each of the three experimental populations of this study (in total *N* = 183 seedlings were used for the three populations).

**Method S2.**
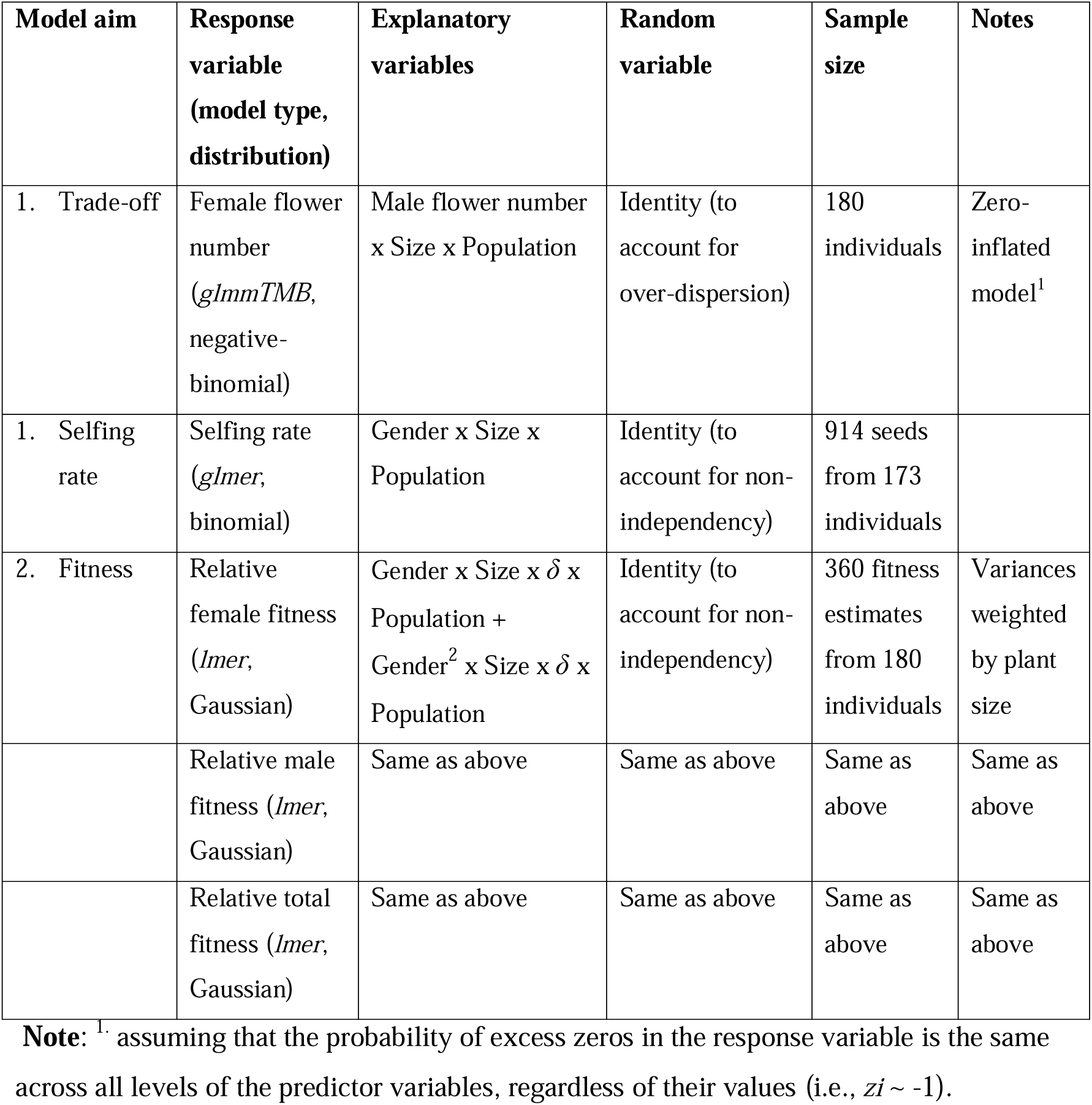
Detailed structure of each regression model used in this study.

**Figure S1.**
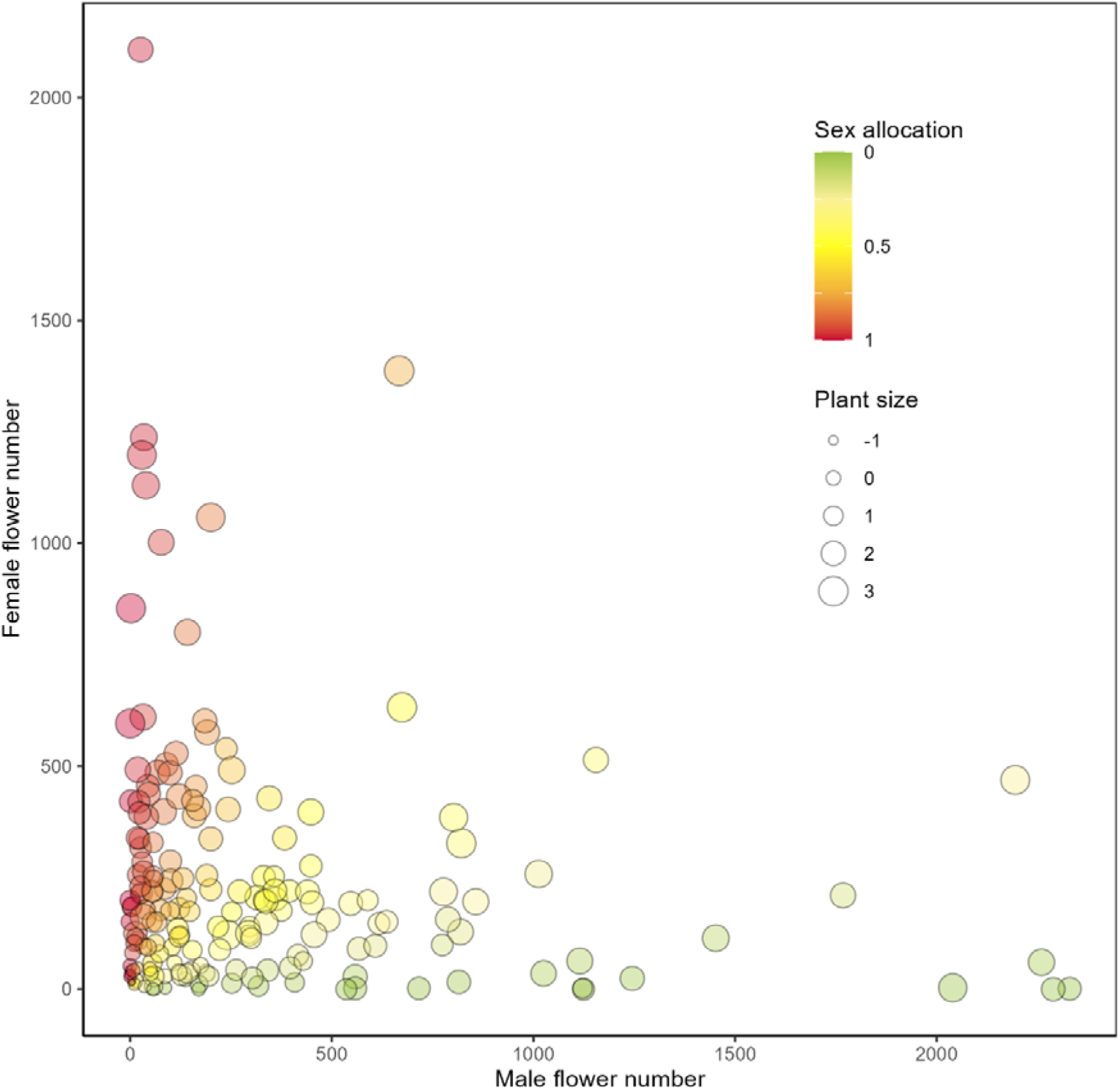
Variation of plant size and sex allocation in the study populations of *Mercurialis annua* (*N* = 180). Actual numbers of male and female flowers of each individual are shown with their sex allocation (maleness). The size of the points indicates the size of the plant. See the main text for details on the calculation of sex allocation and plant size.

**Figure S2.**
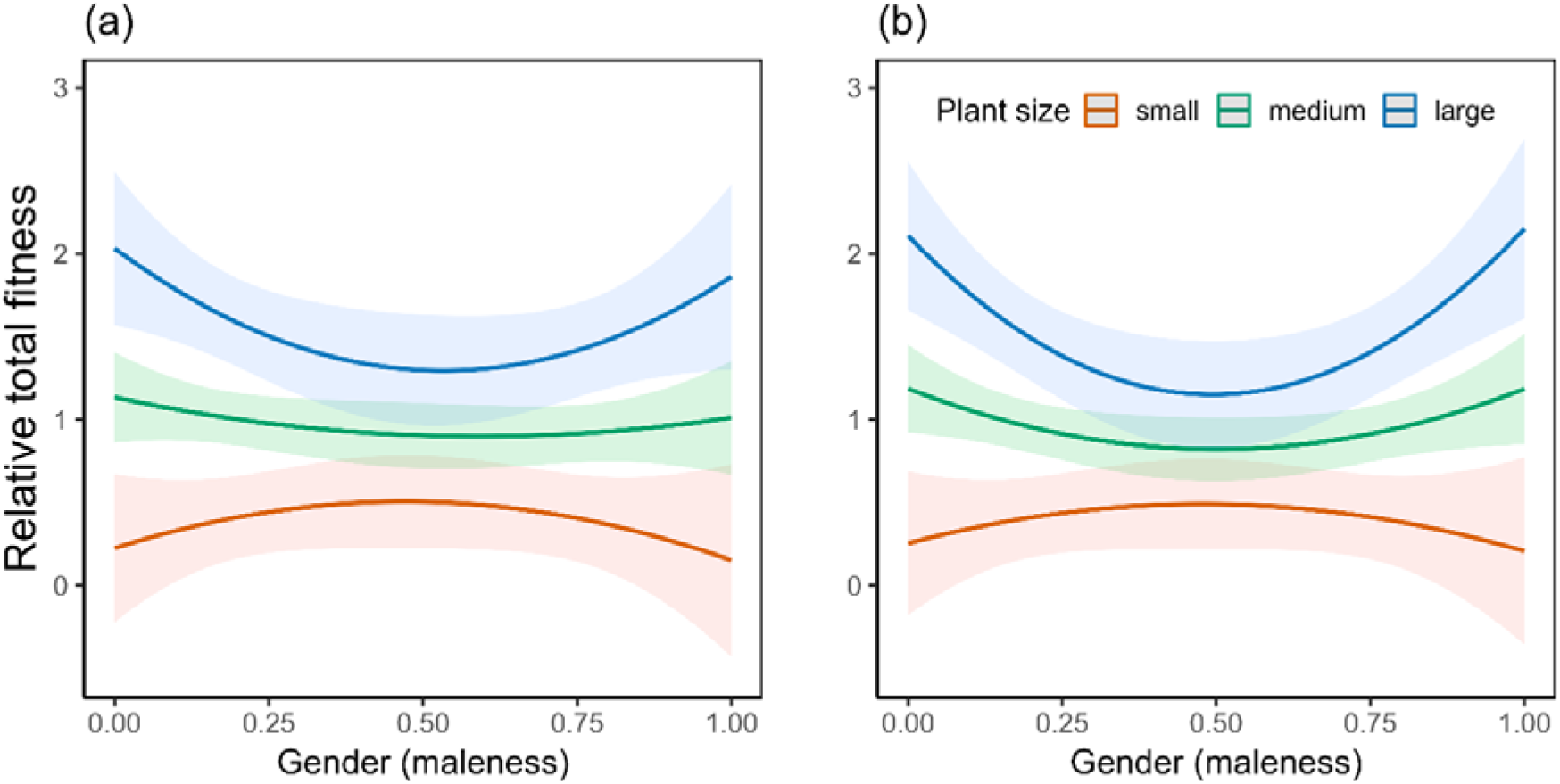
Plots showing the patterns of selection on gender via total fitness under two intermediate levels of inbreeding depression in plants of different sizes. (**a**) When inbreeding depression was 0.2, significant disruptive selection on gender was inferred for plants of large size (quadratic coefficient = 2.59 ± 1.25; *P* < 0.05), whereas no pattern of selection was inferred for plants of medium and small sizes (*P* > 0.05 for both linear and quadratic coefficients). (**b**) When inbreeding depression was 0.7, significant disruptive selection on gender was inferred for plants of medium and large sizes (quadratic coefficient = 1.43 ± 0.71 and 3.92 ± 1.22 with *P* < 0.05 and < 0.01, for medium and large plants, respectively), whereas no pattern of selection was inferred for plants of small size (*P* > 0.05 for both linear and quadratic coefficients). The shaded ribbons indicate the 95% confidence interval of the corresponding regression lines.

**Figure S3.**
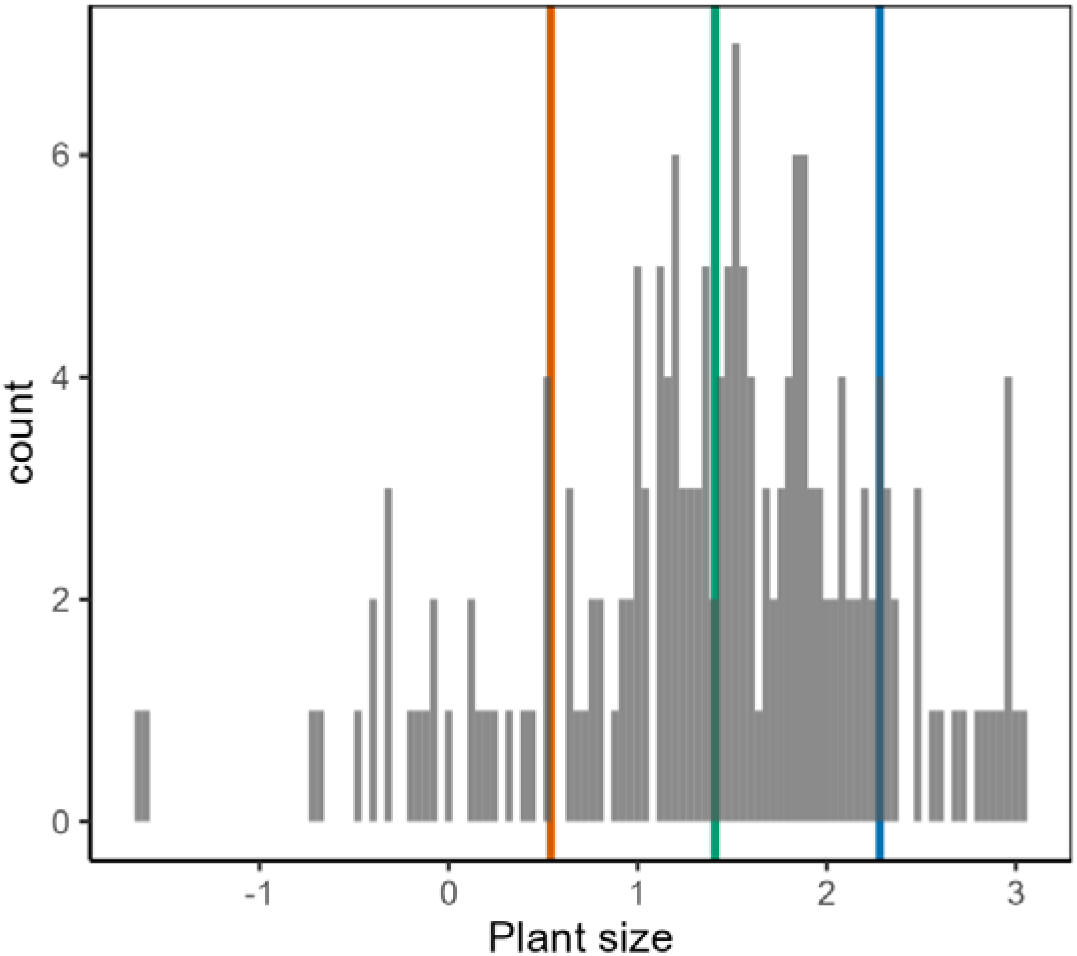
Histogram showing the size of plants in the three experimental populations of *Mercurialis annua* (*N* = 180). The mean size minus SD (at the 16% quantile; hereafter small plants), mean size (at the 50% quantile; hereafter medium plants), and mean size plus SD (at the 84% quantile; hereafter large plants) are indicated by orange, green, and blue vertical lines. Coefficients of an explanatory factor at the three levels of plant size were used to present the significant interactive effect of that factor with plant size on a response variable.

**Figure S4.**
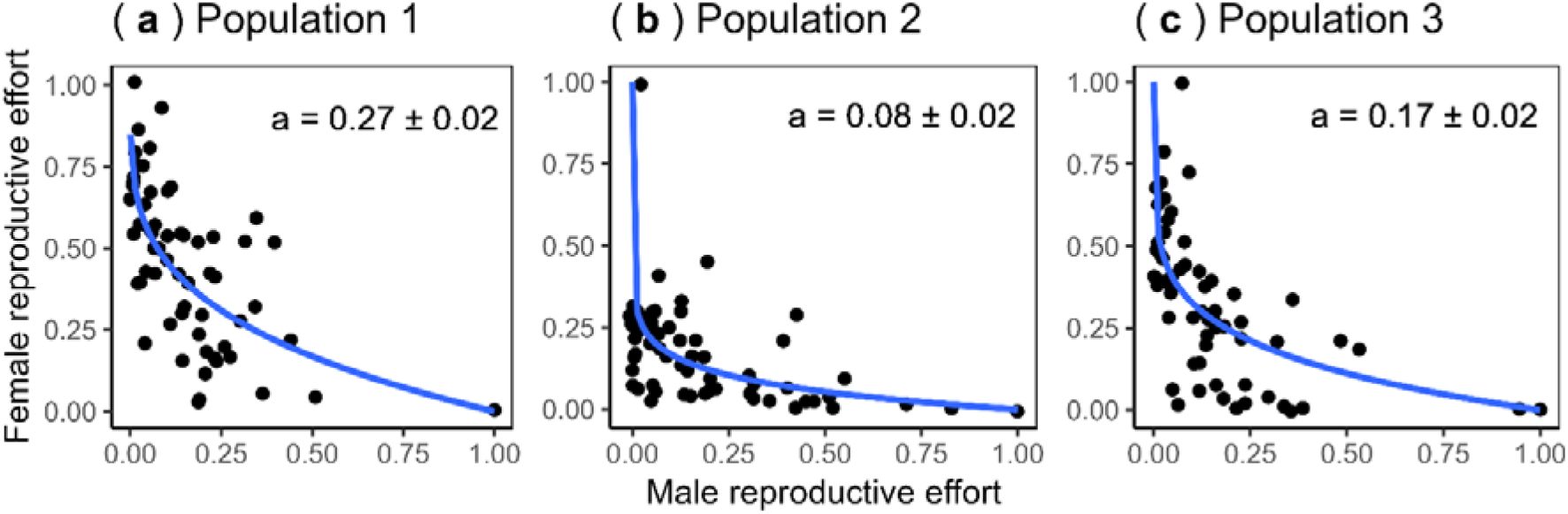
Plots showing the non-linear trade-off curves between female and male functions in studied populations. The non-linear curves were evaluated using non-linear least square regression (*nls* function in *nlme* package; Pinheiro et al., 2022) with the formula below. RE_f_ =1 - RE_m_^a^, where RE and RE_m_ is the female and male reproductive efforts, respectively, defined as the number of flowers of that sex divided by the above-ground biomass of the plant and then relative to the plant with the highest effort in the population. The exponent *a* depicts the nonlinearity of the curves with *a* < 1, = 1, and > 1, indicating inward, linear, and outward trade-offs, respectively. The trade-off curves in the three populations were all concave, with the exponent *a* significantly smaller than one.

**Figure S5.**
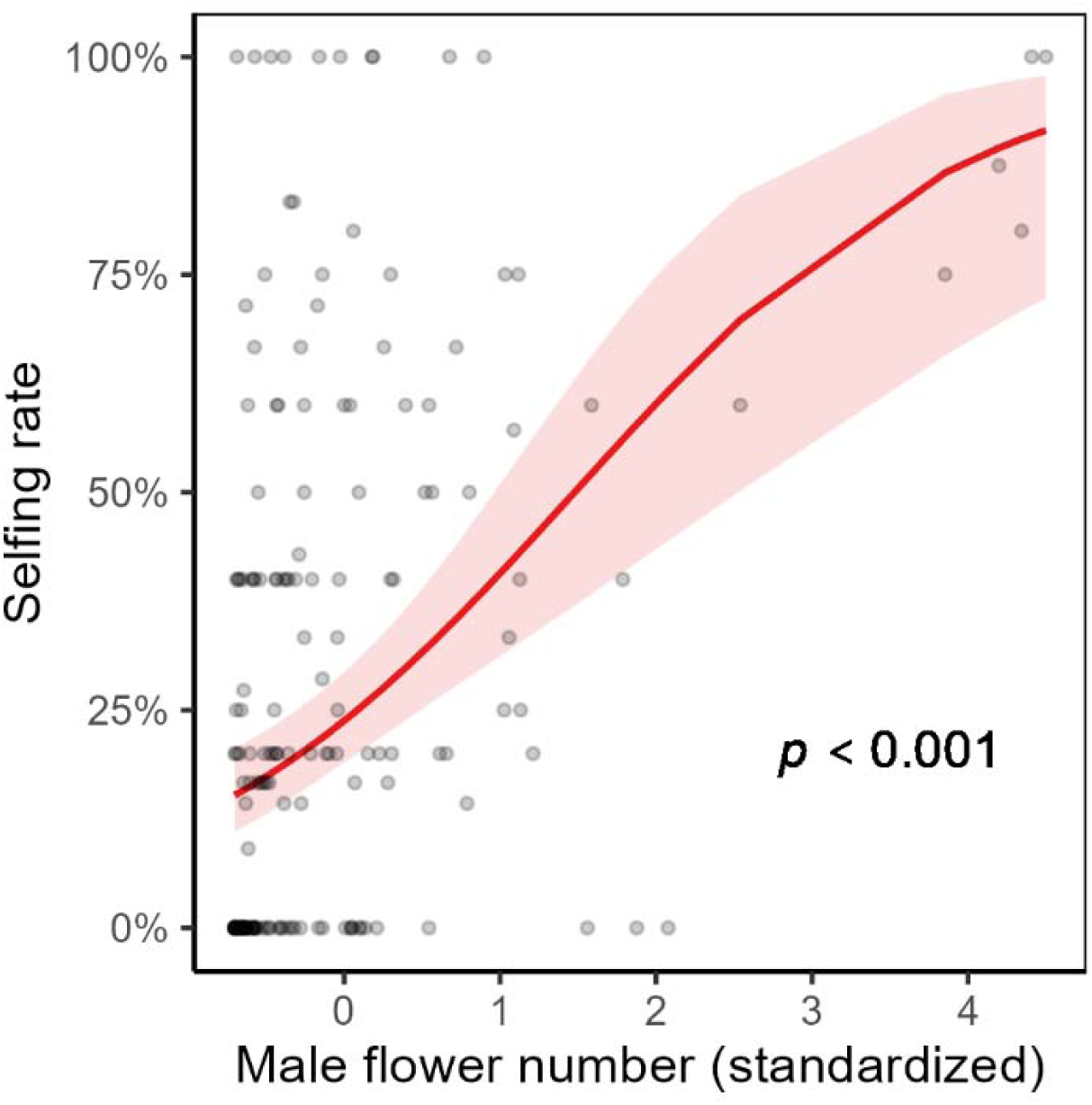
Plot showing the effect of the number of male flowers on the selfing rate in studied individuals (*N* = 172, individuals producing no mature seed were excluded). We used a generalized linear mixed model with a similar structure to the one presented in the main text (see Method S1) except that we replaced the explanatory variables of gender and size with male flower number here. The shaded ribbon indicates the 95% confidence interval of the regression lines.

**Figure S6.**
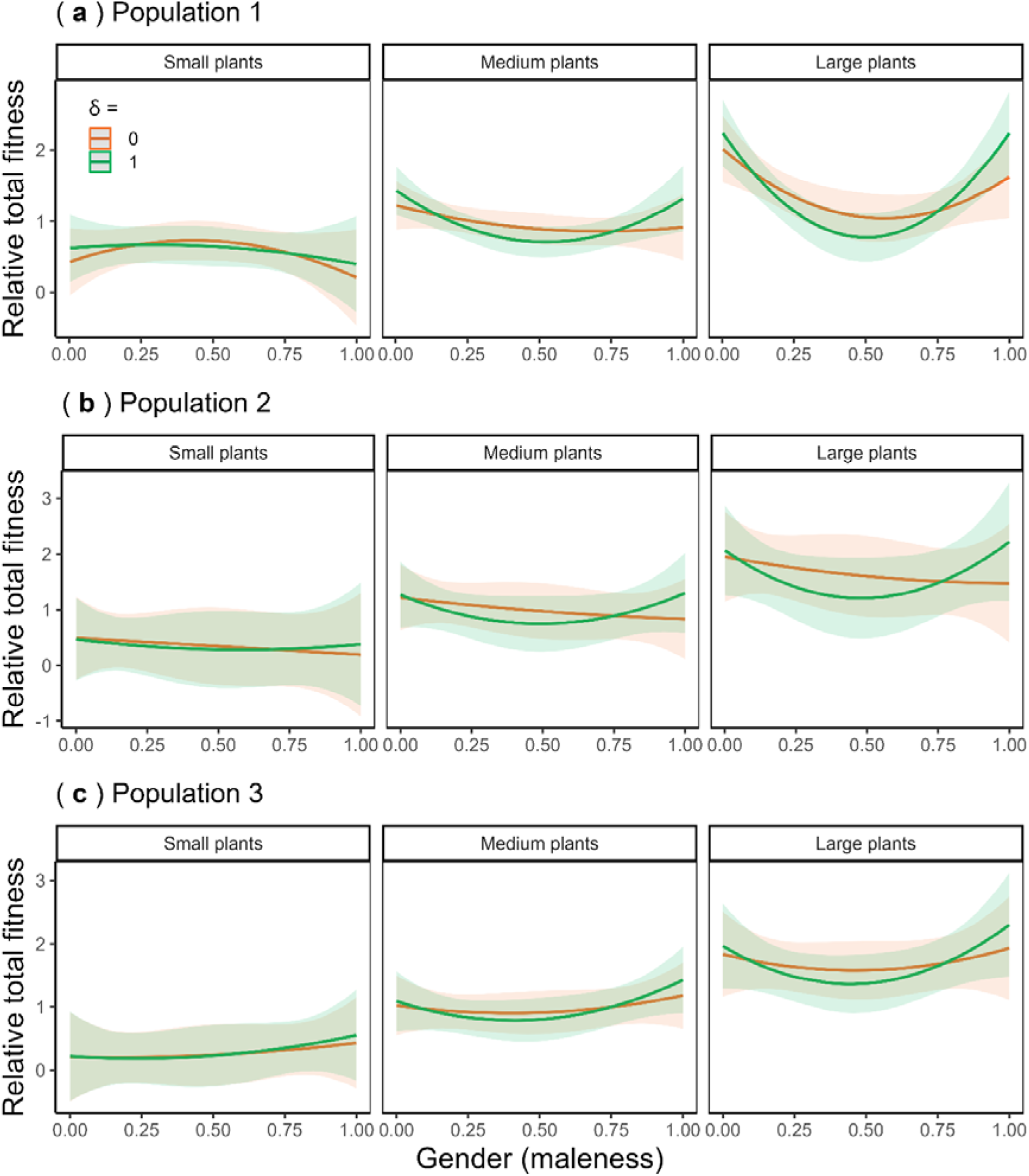
Plots showing the interactive effects of plant size, degree of inbreeding depression, and gender on relative total fitness of *Mercurialis annua* in three experimental populations analysed by three separated linear mixed models. Fitness was estimated under two scenarios of inbreeding depression (δ) of zero and one, depicted by orange and green lines, respectively. The shaded ribbons indicate the 95% confidence interval of the corresponding regression lines. See also Fig. **S7** for the effect sizes.

**Figure S7.**
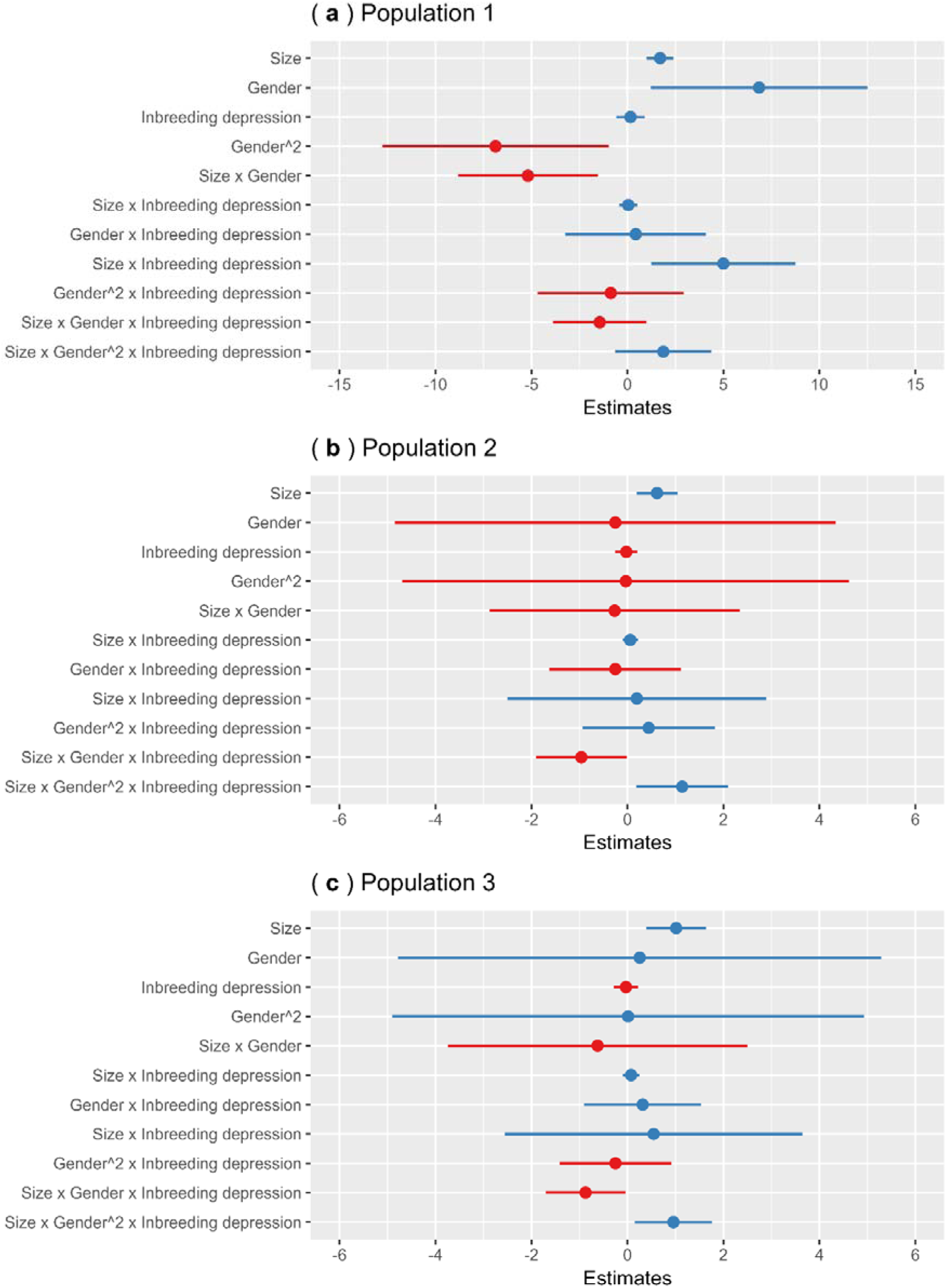
The effect sizes of explanatory factors on relative total fitness of *Mercurialis annua* in three experimental populations analysed by three separated linear mixed models (Fig. **S6**). Positive and negative effects are indicated by blue and red points, respectively.

**Figure S8.**
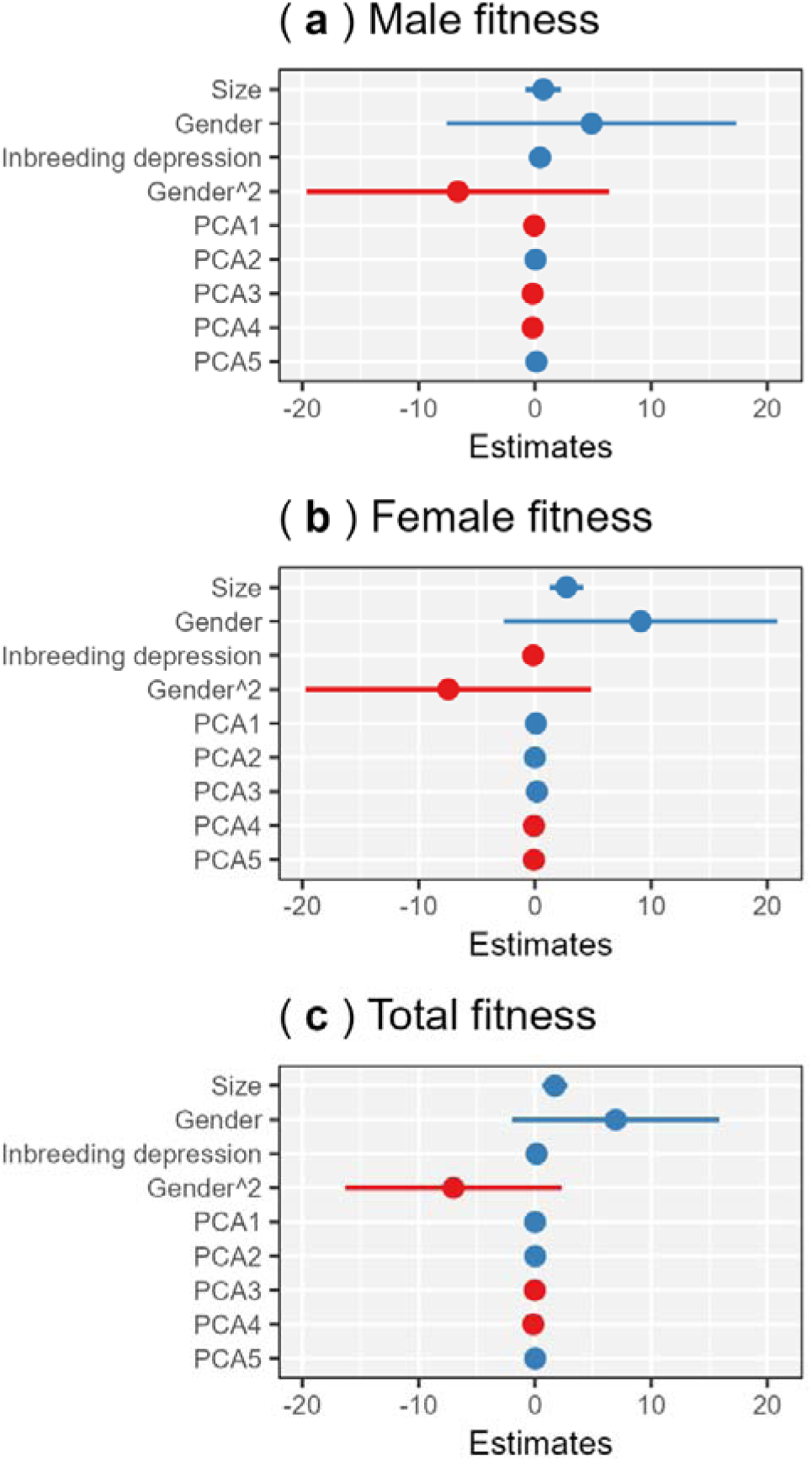
The effect sizes of eight ancillary traits (compressed into 5 PCA axes; see Table **S1**), size, gender (linear and quadratic terms), and inbreeding depression on relative male (**a**), female (**b**), and total (**c**) fitness estimated by linear mixed models (*N* = 175; five individuals with incomplete measurements of traits were not included). The PCA axes were added as single terms into the linear mixed models used for the analysis presented in the main text (see Method **S2** for their structures). For simplicity, only the effect sizes of a selected set of single terms were presented in the figures. The ancillary traits were thought to influence individual fitness via amelioration of pollen dispersal and/or pollen receipt. Nonetheless, according to the analyses, the ancillary traits likely played a minor role in determining individual fitness compared to sex allocation and size.

**Table S1.** Rotated factor matrix for PCA (principal component analysis) on eight ancillary traits and its correlation with gender (male sex allocation) in the individuals of *M. annua* used in this study. The eight ancillary traits were the number of lateral branches, total length of lateral branches, mean length of lateral branches, number of leaves, total leaf area, mean leaf area, leaf dry biomass, leaf specific area sampled from the top 15 cm part of each individual (*N* = 175; 5 individuals with incomplete measurements of traits were not included). PCA axes showed no or weak (i.e., PCA2) correlation with sex allocation, indicating little or no sexual dimorphism in the study populations. See also Fig. **S8** for the effects of the ancillary traits on male, female, and total fitness.

|  | **Mean**<br>**(SD)** | **PCA1** | **PCA2** | **PCA3** | **PCA4** | **PCA5** |
| --- | --- | --- | --- | --- | --- | --- |
| Number of lateral branches | 5.7 (2.5) | -0.10 | -0.58 | 0.16 | -0.64 | 0.05 |
| Total length of lateral branches | 21.2 (13.0) cm | -0.38 | -0.40 | -0.32 | -0.11 | 0.33 |
| Mean length of lateral branches | 3.82 (2.19) cm | -0.36 | -0.05 | -0.52 | 0.40 | 0.33 |
| Number of leaves | 28.1 (14.6) | -0.38 | -0.20 | 0.51 | 0.30 | -0.01 |
| Total leaf area | 47.3 (34.2) cm<sup>2</sup> | -0.50 | 0.21 | 0.20 | -0.04 | -0.17 |
| Mean leaf area | 1.82 (1.34) cm<sup>2</sup> | -0.23 | 0.45 | -0.39 | -0.52 | -0.20 |
| Leaf dry biomass | 160 (107) mg | -0.51 | 0.10 | 0.12 | 0.02 | -0.37 |
| Leaf specific area | 0.29 (0.06) cm<sup>2</sup>mg<sup>-1</sup> | -0.10 | 0.44 | 0.37 | -0.22 | 0.76 |
| Cumulative variance explained (%) |  | 0.41 | 0.63 | 0.79 | 0.89 | 0.98 |
| Correlation coefficients with sex allocation ($p$-value) | | 0.07 (n.s.) | -0.21 ($p < 0.001$) | 0.001 (n.s.) | 0.02 (n.s.) | 0.05 (n.s.) |

**Table S2.**
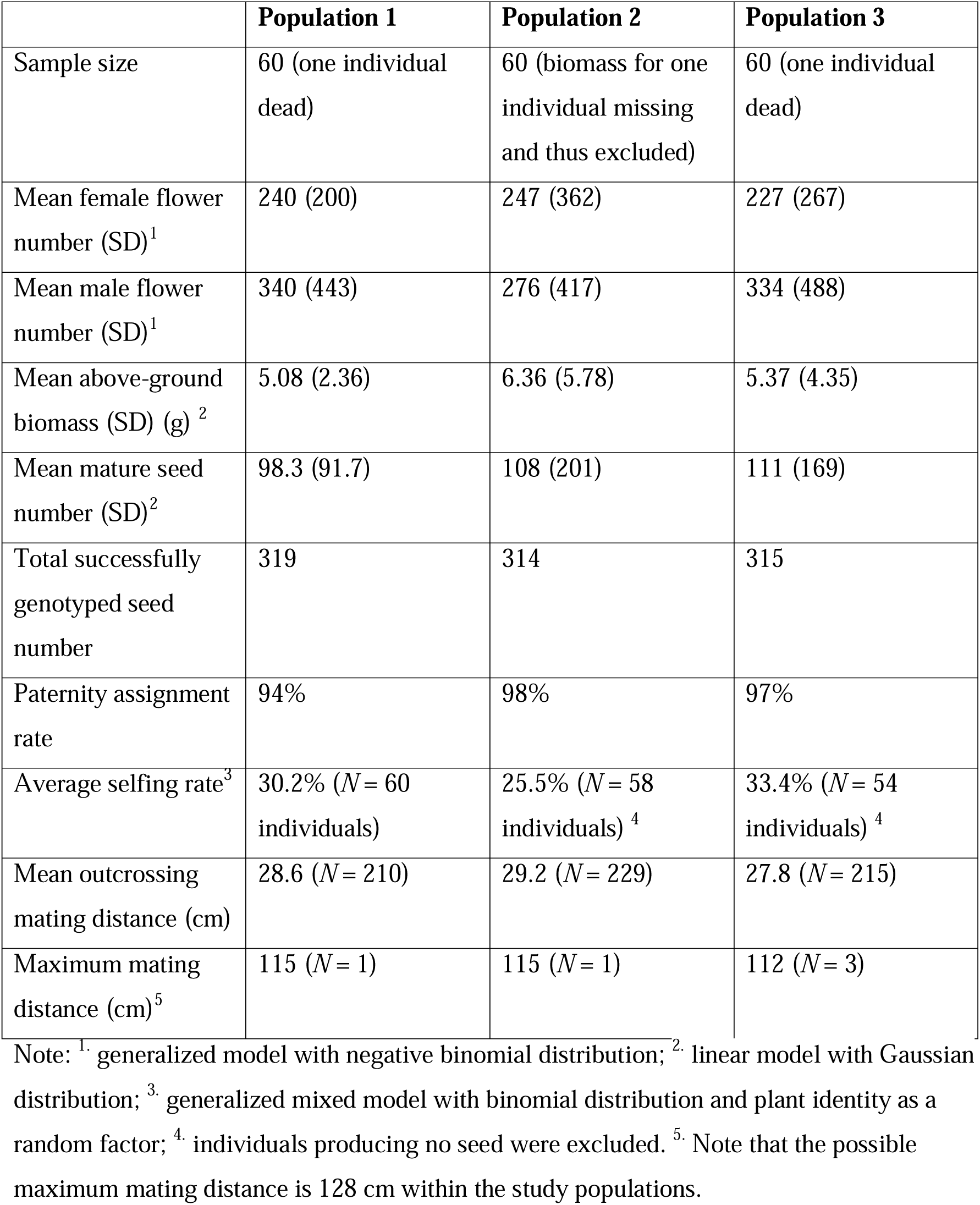
Details of sex allocation, biomass, and paternity analyses of the three experimental populations. Population showed no difference in all the variables tested (*P* > 0.05; see the note below).

**Table S3.**
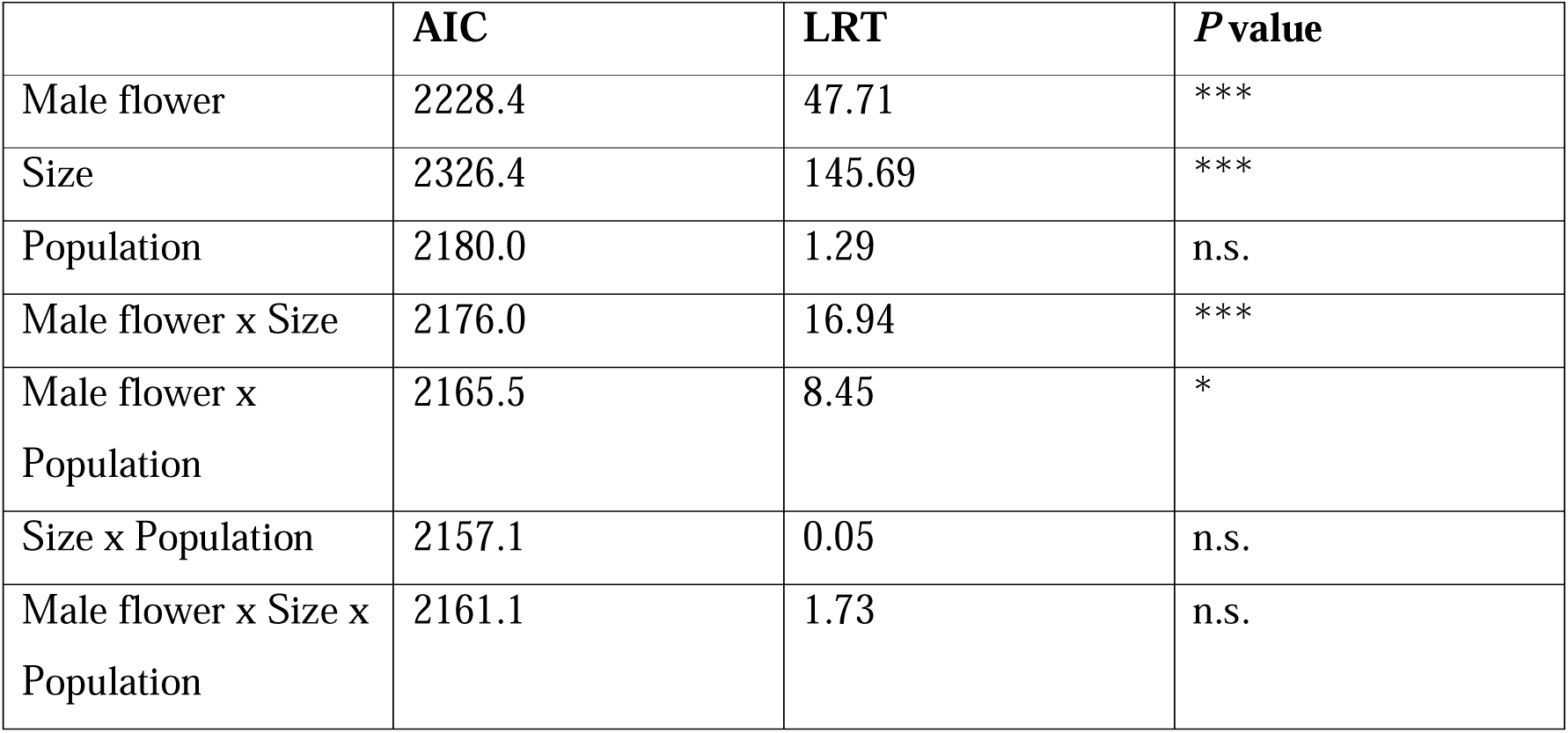
Summary table of the general effects of male flower number, size, and population on female flower number estimated by a generalized linear mixed model. The *P* values were extracted using likelihood ratio tests. Notes: n.s. *P* > 0.05, ∗ *P* < 0.05, ∗∗ *P* < 0.01, ∗∗∗ *P* < 0.001.

**Table S4.**
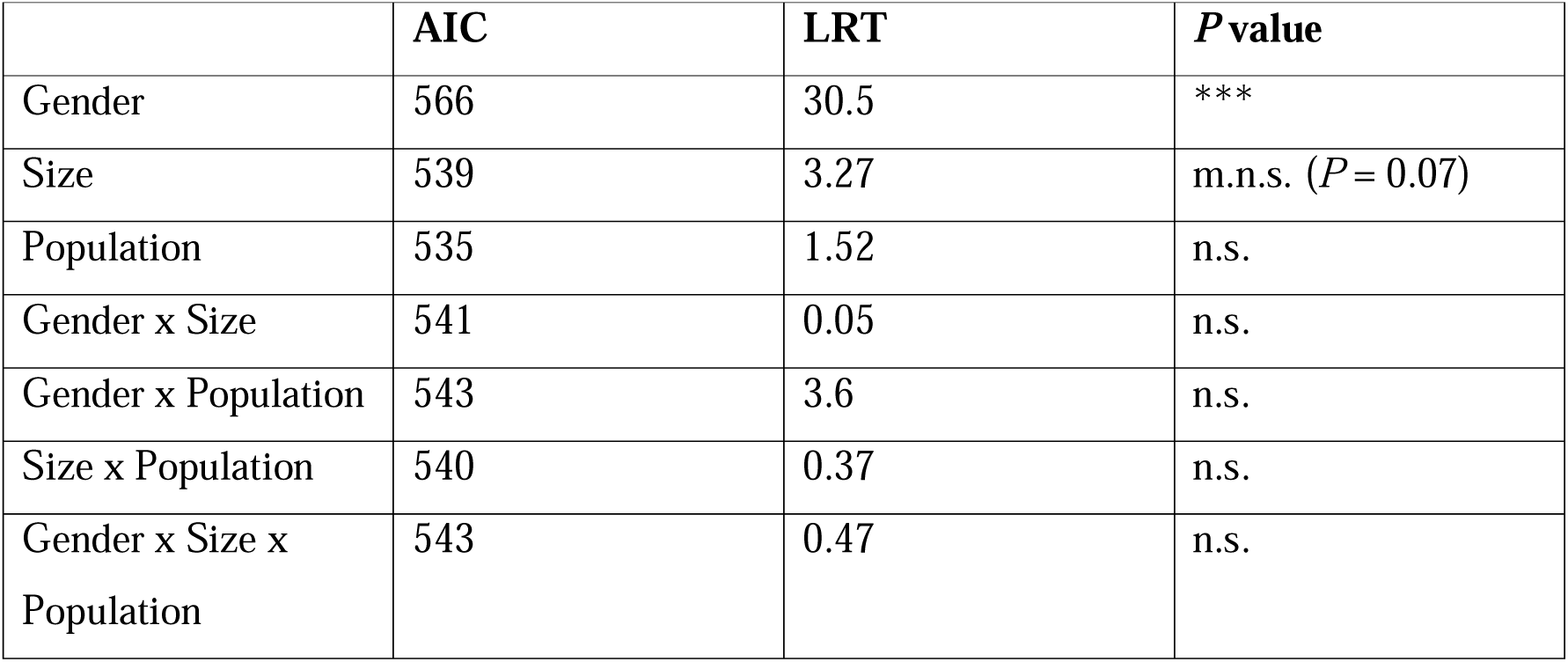
Summary table of the general effects of size, gender, and population on the selfing rate estimated by a generalized linear mixed model. The *P* values were extracted using likelihood ratio tests. Notes: n.s. (non-significant) *P* > 0.1, m.n.s. (marginally non-significant) *P* < 0.1, ∗ *P* < 0.05,∗∗ *P* < 0.01, ∗∗∗ *P* < 0.001.

**Table S5.**
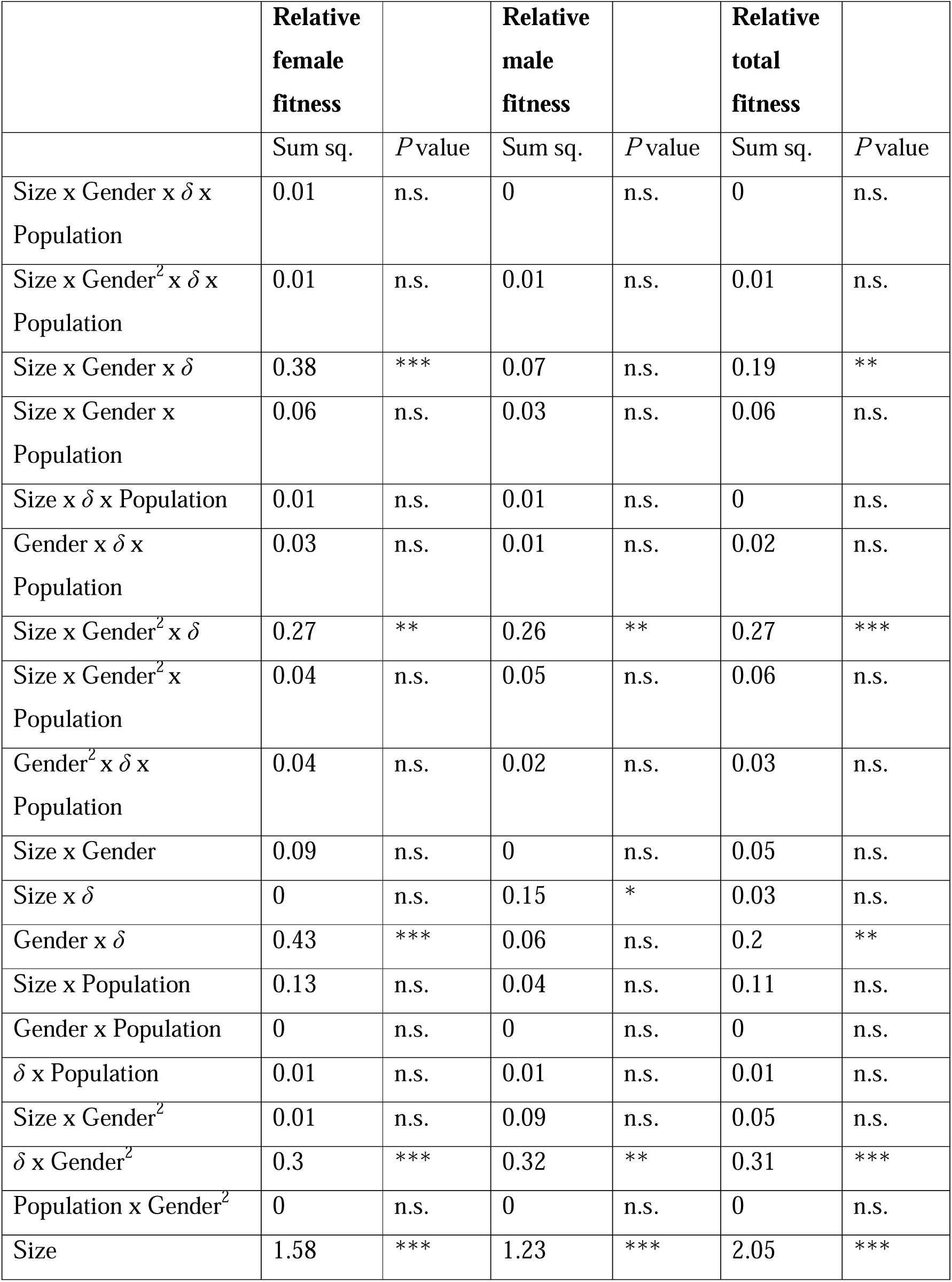
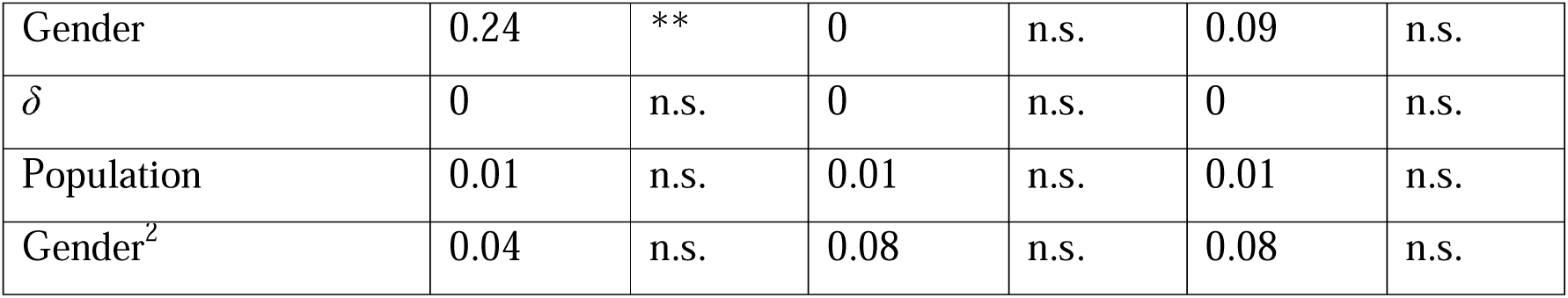
Summary table of the general effects of size, gender (linear and quadratic terms), scenarios of inbreeding depression (δ), and population on relative female, male, and total fitness estimated by linear mixed models. Notes: n.s. *P* > 0.05, ∗ *P* < 0.05, ∗∗ *P* < 0.01, ∗∗∗ *P* < 0.001.

